# Compensatory aortic remodeling in Marfan syndrome protects against sexually dimorphic rupture during a BAPN challenge

**DOI:** 10.1101/2022.06.21.497029

**Authors:** D. Weiss, B.V. Rego, C. Cavinato, D.S. Li, Y. Kawamura, N. Emuna, J.D. Humphrey

## Abstract

Transmural rupture of the aorta is responsible for significant morbidity and mortality; it occurs when wall stress exceeds local wall strength. Amongst other conditions, the aortic root and ascending aorta become vulnerable to dissection and rupture in Marfan syndrome, a connective tissue disorder that results in a progressive fragmentation and degradation of the elastic fibers of the aortic wall. Whereas competent elastic fibers are critical for aortic functionality, cross-linked collagen fibers endow the aorta with its stiffness and strength. In this paper, we contrast progressive degeneration of the ascending aorta in male and female Marfan and wild-type mice, with and without chronic exposure to a potent inhibitor of lysyl oxidase (β-aminopropionitrile, or BAPN), to examine effects of extracellular matrix cross-linking in aortic dilatation and rupture. We found a strong sexual dimorphism in aortic dilatation in Marfan mice and aortic rupture in wild-type mice, but also a compensatory remodeling of the aorta that protected the Marfan aorta against lethal rupture despite a strong BAPN challenge. This compensation appears to be mediated via increased lysyl oxidase in the female and especially male Marfan aorta, resulting in improved collagen fiber stability and integrity, particularly of fibril bundles in the adventitia.

## INTRODUCTION

Marfan syndrome (MFS) is a connective tissue disorder that results from mutations in the gene (*FBN1*) that encodes fibrillin-1^1^, an elastin-associated glycoprotein that plays multiple roles in extracellular matrix development and homeostasis^2^, including contributions to the long-term stability of normal elastic fibers. Although diverse tissues are affected, premature mortality in MFS patients results from rupture of the thoracic aorta, often secondary to aneurysmal dilatation. Much has been learned about this aortic condition from clinical studies that include genetics, cell biology, histopathology, *in vivo* imaging, and computational modeling, but mouse models continue to provide valuable insight. Two commonly used models of MFS are *Fbn1*^*mgR/mgR*^ mice, which are hypomorphic for wild-type fibrillin-1^3^, and *Fbn1*^*C1041G/+*^ mice (a.k.a. *Fbn1*^*C1039G/+*^), which express a mutant form of fibrillin-1^4^. Both models have been studied biologically and biomechanically, with reports of increased pulse wave velocity and abnormal active and passive biomechanical properties^5–10^. In particular, the biomechanical phenotype of the aneurysmal MFS aorta is characterized by decreased elastic energy storage, consistent with compromised elastic fiber integrity, and increased circumferential material stiffness, consistent with an altered collagen fiber architecture that results from dysfunctional mechano-sensing and/or mechano-regulation by the cells^11,12^.

Notwithstanding the ubiquitous fragmentation and degradation of elastic lamellae in the aorta of MFS patients and mouse models, which drives dilatation^7,13^, rupture of the aorta depends in large part on a loss of integrity of the fibrillar collagens^14–16^, which normally endow the aorta with both stiffness and strength. Recent RNA sequencing revealed increased transcripts for fibrillar collagens in MFS in humans and both common mouse models^17–19^, consistent with histological evidence of accumulated collagen. Whereas compromised elastic fiber integrity can have direct effects on macroscale collagen properties, including reduced collagen undulation and thus increased material stiffness^20^, fibrillar collagens near and in the adventitia also exhibit altered microstructure and micromechanical properties in MFS^21,22^. That is, despite the monogenic mutation affecting fibrillin-1, there appears to be important effects on collagen fibrillogenesis and organization, which involve associated matrix constituents (e.g., biglycan^23^) and require intra- and inter-molecular cross-linking to yield mechanically competent collagen fibers. There is, therefore, a pressing need to assess better the state of collagen in MFS, not just the elastic fibers.

Lysyl oxidase is a copper-dependent enzyme that initiates aldehyde-based covalent cross-linking of newly synthesized elastic and collagen fibers, hence contributing significantly to the biomechanical properties of these fundamental structural proteins of the aorta^24,25^. Importantly, mutations in the gene (*LOX*) that encodes lysyl oxidase predispose patients to thoracic aortic aneurysms and dissections^26^, and *Lox* mutant mice exhibit similar aortopathies^27^. A recent study using both common mouse models of MFS^28^ suggested that enhanced LOX-mediated collagen accumulation helps to protect against aortic dilatation (*Fbn1*^*C1041G/+*^ mice) and rupture (*Fbn1*^*mgR/mgR*^ mice), the former consistent with a prior study that showed attenuated aortic dilatation in *Fbn1*^*C1041G/+*^ mice when antagonizing miR29b, a microRNA that slows collagen production^29^. There was, however, no detailed assessment of the biomechanical phenotype of the aorta in any of these mouse studies. Although cell biological and histological findings can be highly suggestive, there is a need for direct functional readouts of associated consequences on the mechanical state of the aorta because it is mechanical failure – dissection and rupture – that is ultimately responsible for the morbidity and mortality. In this paper, we quantify consequences of LOX inhibition in the *Fbn1*^*C1041G/+*^ mouse model of MFS using both standard and novel multi-modality biaxial quantification of mechanical properties with associated intramural imaging. To begin to delineate contributions of elastic fibers and fibrillar collagens to early disease progression, we contrast outcomes in female and male mice when LOX inhibition is initiated at 4 weeks of age, after elastic lamellar structures but not intra-lamellar elastic or collagen fibers have matured, and at 8 weeks of age, after the aorta has matured biomechanically^30^.

## RESULTS

### Age and sex are important biological variables in lysyl oxidase mediated aortic integrity

Values of 12 key measured and calculated geometric and mechanical metrics for the ascending aorta are compared in **Tables S1,S2** (mean ± standard error of mean, *n* = 5 per group) for all 16 Groups of interest: male and female wild-type (MWT, FWT) and male and female *Fbn1*^*C1041G/+*^ Marfan (MMFS, FMFS) mice, each at 8 versus 12 weeks of age without or with preceding 4-week exposures to the potent inhibitor of lysyl oxidase (β-aminopropionitrile, or BAPN). Standard pressure-diameter data revealed key differences under physiologic loading (**Fig S1A**). Responses were similar across the 4 basic Groups at 8 weeks of age, though with an emerging MFS phenotype that was greater at 12 weeks of age in male but not female *Fbn1*^*C1041G/+*^ mice. Exposure to BAPN from 4 to 8 weeks of age had modest effects on this characteristic behavior of the WT aortas but induced marked changes in the MFS aortas, especially in males. Exposure to BAPN from 8 to 12 weeks of age resulted in similar, though attenuated results in the MFS mice. Associated wall stress-stretch behaviors were similar (**Fig S1B**). Together, these results reveal clear age- and sex-dependent differences between WT and MFS aortas during a BAPN challenge, thus motivating a careful examination of the intrinsic properties that define the aortic phenotype.

### The MFS phenotype is characterized by progressive decreases in energy storage and increased material stiffness prior to aneurysmal dilatation

The primary biomechanical function of the aorta is to store elastic energy during systole and to use this energy during diastole to recoil the wall and work on the blood to augment flow. Elastic energy storage capacity was lower in the MFS aorta at 8 weeks of age relative to age- and sex-matched WT controls and this difference worsened at 12 weeks of age (**Fig 1B,J**): ~27 and 48% reductions in males and 27 and 31% reductions in females at 8 and 12 weeks in MFS, respectively. These progressive changes were mirrored by modest but progressive decreases in the *in vivo* value of axial stretch (**Fig 1F,N**), again more so in males than in females in MFS: ~6 and 12% reductions in MFS males, ~0 and 5% reductions in MFS females at these two ages. Conversely, there was an early marked and sustained increase in the circumferential material stiffness in MFS (**Fig 1C,K**), again more so in males (~1.95- and 2.05-fold increases at 8 and 12 weeks of age) than in females (1.72- and 1.42-fold increases). Changes in axial material stiffness were less marked at both ages (**Fig 1G,O)**. There was little difference in wall thickness at 8 weeks of age between MFS and WT, but the aortic wall thickened by ~21% in the male MFS mice at 12 weeks (**Fig 1E,M)**, consistent with the greater disease presentation and progression in males. Importantly, these biomechanical changes manifested despite modest aortic enlargement in the MFS mice (8-10% in males, 1-2% in females; **Fig 1A,I**), well less than the definition of an aneurysm (≥50%), thus revealing that that early changes in biomechanical composition and function in MFS precede aneurysmal dilatation.

**Figure 1.**
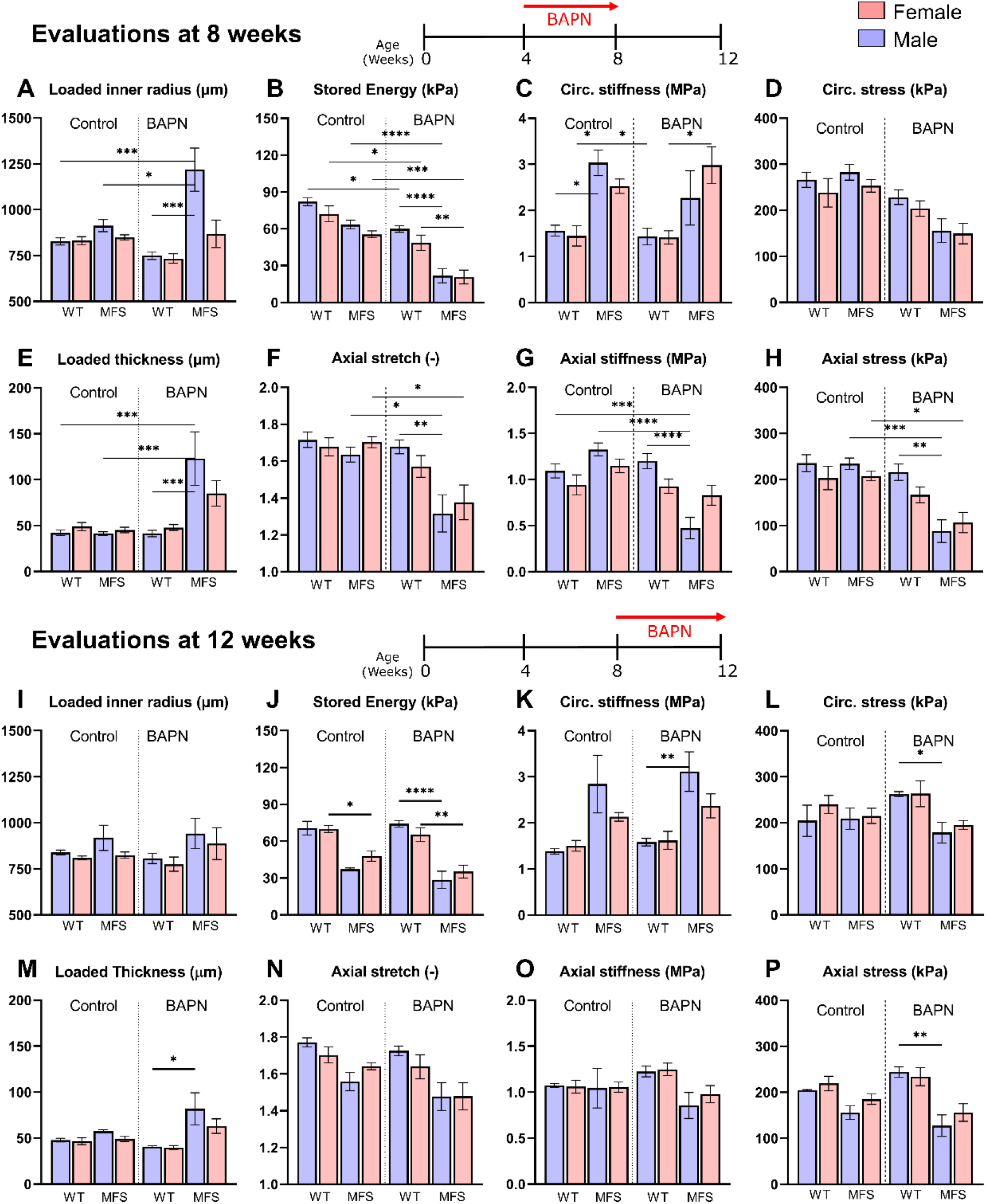
Comparison of 8 key geometric and mechanical metrics (panels A-P) for all 16 Groups: female (F) and male (M), wild-type (WT) and *Fbn1*^*C1041G/+*^ Marfan (MFS) ascending aortas with BAPN given for 4 weeks beginning either at 4 weeks of age (then evaluated at 8 weeks) or at 8 weeks of age (then evaluated at 12 weeks of age), with age- and sex-matched controls not receiving BAPN. Values are means ± standard deviation (*n*=5 per group, hence 80 aortas tested) inferred from *ex vivo* biaxial testing of excised segments under physiological conditions. Statistical comparisons by non-parametric Kruskal Wallis test followed by Dunn’s post-hoc test for multiple comparisons.. \*\*\*\**p*<0.0001, \*\*\**p*<0.001, \*\**p*<0.01, \**p*<0.05. See Figure S1 for pressure-diameter and stress-stretch relations and Tables S1 and S2 for all numerical values.

### BAPN reduces energy storage with modest effects on stiffness in MFS

BAPN only blocks cross-linking of newly deposited elastic and collagen fibers, hence we contrasted effects of 4-week exposures to BAPN starting at either 4 (during aortic development) or 8 (after aortic maturity) weeks of age, with evaluations at 8 and 12 weeks of age consistent with those in the non-treated mice. BAPN markedly reduced energy storage when started at 4 weeks of age in WT (25% reductions in males, 32% in females) and especially in MFS (66% in males, 62% in females) mice; reductions were modest when BAPN was started at 8 weeks of age in both WT (<10% in both sexes) and MFS (22% in males, 27% in females) mice, recalling that energy storage was naturally lower in MFS at 12 weeks of age (**Fig 1B,J**). These trends were paralleled by those for axial stretch (**Fig 1F,N**), both when BAPN was started at 4 weeks of age (<6% reductions in male and female WT, 19% and 18% reductions in male and female MFS) and when started at 8 weeks of age (<5% reductions in male and female WT, 5-10% reductions in male and female MFS). Results for circumferential material stiffness were mixed (**Fig 1C,K**). This stiffness changed less than 3% in male and female WT mice but decreased 25% and increased 17% in male and female MFS mice, respectively, when BAPN was started at 4 weeks of age. By contrast, this stiffness increased ~10% in male and female WT and MFS mice when BAPN was started at 12 weeks of age. Results were similar in the axial direction, though less dramatic (**Fig 1G,O**). Hence, BAPN reduced circumferential stiffness only in the male MFS mice, and only when BAPN was started early (at 4 weeks of age).

### BAPN increases wall thickness, and decreases wall stresses, in MFS but not WT mice

One of the most dramatic changes seen in any of the geometric and mechanical metrics (**Tables S1,S2**) was in aortic wall thickness in the MFS mice when BAPN was started during development (**Fig 1E,M**). In particular, wall thickness changed less than 3% in male and female WT mice when BAPN was started at 4 weeks of age while it decreased ~15% in WT mice when started at 8 weeks of age. By contrast, wall thickness increased dramatically in MFS mice, especially when BAPN was started at 4 weeks of age (~3-fold increases in male and ~1.9-fold in female mice) but also when started at 8 weeks of age (41% increases in male and 28% in female mice). These changes in wall thickness combined with the aforementioned changes in circumferential stiffness similarly affected both distensibility and calculated circumferential structural stiffness (**Fig S2**). Standard histological examinations of fixed aortic cross-sections stained with Movat pentachrome revealed that aortic thickening in MFS in response to BAPN arose from complex changes in mural content, mainly increases in glycosaminoglycans (GAGs) and adventitial collagens, especially when BAPN was started earlier, but without any consistent grossly adverse effects of BAPN on the elastic lamellae (**Figs S3-S5**). Importantly, the increase in adventitial thickness in the MFS aortas was accompanied by an increase in adventitial cell density, which was in contrast to a loss of medial smooth muscle cells and endothelial cell density (**Fig 2A**). Although the Movat staining did not reveal many changes in elastic fiber area or percent, two photon fluorescence imaging of non-fixed, physiologically loaded segments revealed increases in elastin porosity and pore size in the young male MFS aortas following BAPN-exposure, particularly in the inner and outer portions of the media (**Fig 2B-D, Fig S6**). Importantly, increased inner diameter correlated strongly with increased porosity (**Fig 2E**).

**Figure 2.**
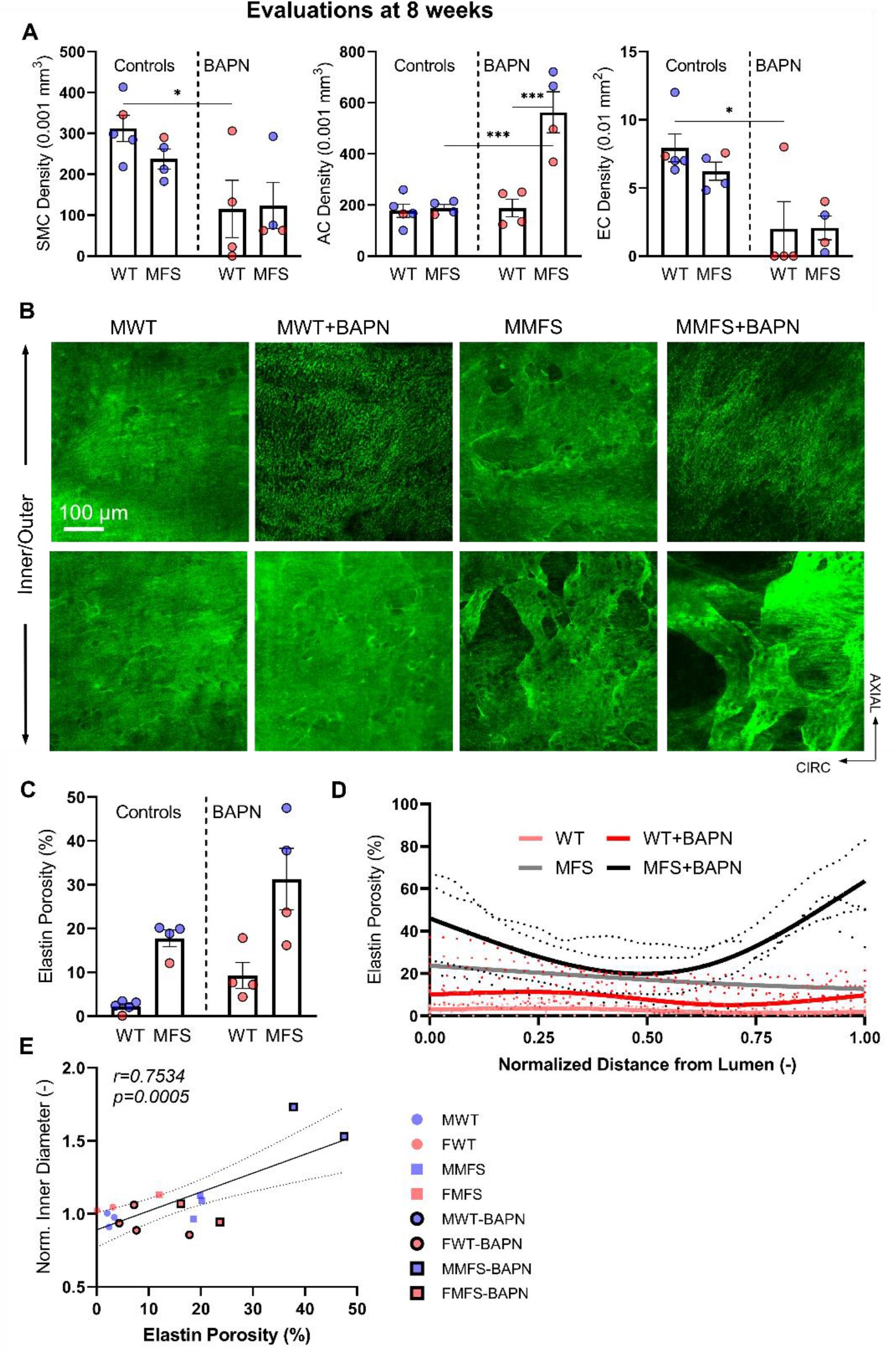
A. Multiphoton microscopy revealed in a sub-set of vessels evaluated at 8 weeks of age that there was a general decrease in endothelial and smooth muscle cell density in WT and MFS aortas following BAPN exposure, but a marked increase in adventitial cell density in BAPN exposed MFS aortas, particularly in males. B, C. Two photon fluorescence revealed further that elastin porosity (and also pore size - **Fig S6**) were both initially greater in MFS than WT aortas, and these measures of compromised elastin increased in MFS aortas following early BAPN exposure. Importantly, the latter tended to manifest primarily in the inner and outer media, not the middle portion of the media. See **Figs S2, S3** for detailed histological quantification based on fixed, Movat-stained cross-sections

Many tacitly assume that wall stress is higher in dilated and aneurysmal arteries, but such stress results from a complex combination of changes in inner radius, wall thickness, and axial stretch under physiological biaxial loading. Importantly, noting that tail-cuff measured systolic blood pressures did not differ across Groups, values of wall stress were similar in male and female WT and MFS mice at both 8 weeks (with circumferential values ~240-280 kPa) and 12 weeks (slightly lower at ~205-240 kPa) of age (**Fig 1D,H**). There was, however, a 9-18% decrease in biaxial wall stress in the male and female WT mice when BAPN was started at 4 weeks of age and a 6-27% increase in the WT mice when BAPN was started at 8 weeks of age. By contrast (**Fig 1L,P**), consistent with the marked changes in wall thickness, there was a dramatic decrease in wall stress in the MFS mice when BAPN was started at 4 weeks of age (reductions of 45-63% in males and 41-49% in female) and a similar though less dramatic decrease when BAPN was started at 8 weeks of age (9-18%, slightly greater in males).

### Multi-modality imaging data confirm standard biaxial results, but emphasize local variations

Gross examinations revealed spatial heterogeneity in the aortic phenotype, especially in the young male MFS aortas wherein particular regions tended to have greater dilatation (**Fig 3A**). Hence, given the overall more dramatic effects of BAPN when administered during development, the aforementioned bulk biomechanical metrics were evaluated independently, in a sub-set of 8-week-old male and female WT and MFS aortas with and without BAPN exposure (*n*=23, 12 male and 11 female), by combining panoramic digital image correlation (pDIC) + optical coherence tomography (OCT) to measure local geometric metrics and to estimate local material properties under physiological biaxial loading. These results confirmed the findings from standard biaxial testing (cf. **Fig 1**) but quantified the heterogeneous biomechanical changes around the circumference and along the length of the ascending aorta (**Fig 3B,C, Figs S7-10**). Again, values of circumferential material stiffness were higher in MFS compared to age- and sex-matched controls, and were not affected much by BAPN. The pDIC + OCT data also quantified marked regional variations in wall thickness, particularly in the male MFS aorta. Importantly, these regions of reduced thickness were observed in vessels that did not rupture *in vivo*.

**Figure 3.**
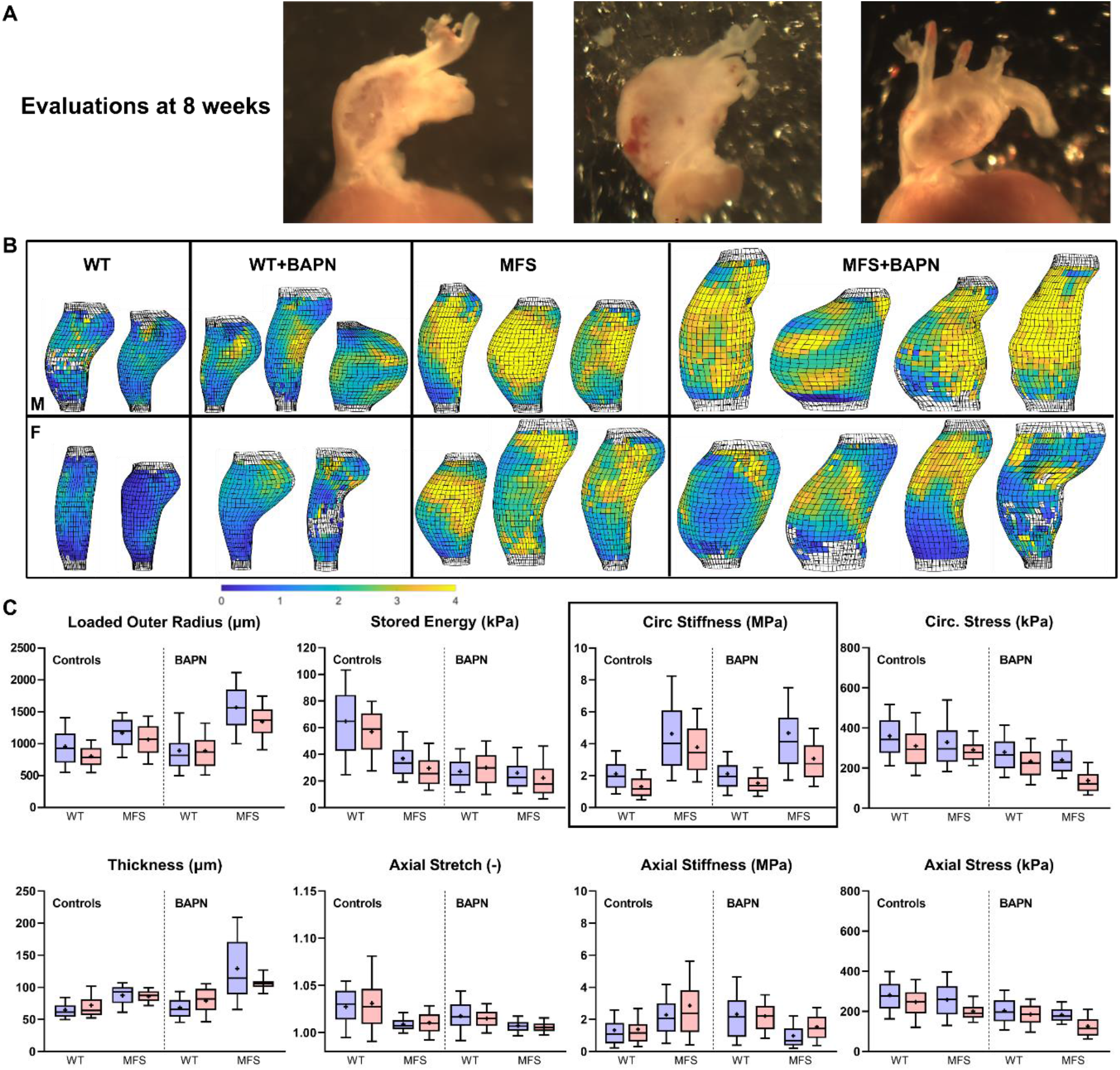
A. Illustrative gross images of three male MFS aortas showing marked regional thinning of the wall, noting that these vessels did not rupture *in vivo*. B, C. Multimodal (pDIC + OCT) *ex vivo* testing under physiological loading confirmed, in a sub-set of 8-week old mice (*n*= 23 samples), findings from standard biaxial testing, but further revealed regional heterogeneities in most geometric and mechanical metrics (see **Figs S7-S10** for sample-specific distributions). Highlighted for illustrative purposes is the, perhaps surprising, lack of a dramatic effect of *in vivo* BAPN exposure on circumferential material stiffness.

### BAPN results in lethal aortic rupture in young WT mice, especially males, but less so in MFS mice

Although the most dramatic changes in standard geometric and mechanical metrics for the aorta (**Tables S1,S2**) occurred in the male MFS mice when BAPN was started at 4 weeks of age (especially increased luminal radius and wall thickness, decreased axial stretch and energy storage, and decreased biaxial stress), the most dramatic effect on aortic rupture and thus mortality was in male WT mice when BAPN was started at 4 weeks of age. That is, whereas all mice survived to the intended endpoint when BAPN was started at 8 weeks of age, only 41/74 (55%) of the WT mice started on BAPN at 4 weeks of age survived while 80/87 (92%) of the MFS mice started on BAPN at 4 weeks survived (**Fig 4A**). Importantly, of the 41 surviving younger WT mice, 33 (80%) were female and only 8 (20%) were male. Given that material failure occurs when wall stress exceeds wall strength, recall that exposure to BAPN from 4 to 8 weeks of age resulted in a modest reduction in circumferential stress in WT mice (~14% lower) and a more dramatic reduction in MFS mice (~43% lower). That wall stress was lower in MFS than WT aortas is consistent with the greater rupture in WT, but the aortic stresses were yet lower than normal in all BAPN exposed mice, thus suggesting compromised wall strength. Pressure-burst tests confirmed significantly lower burst pressures for male WT aortas and a trend toward lower burst pressures for female WT aortas when BAPN was started at 4 weeks of age. Conversely, burst pressures were higher, not lower, in aortas from male and female MFS mice when BAPN was started at 4 weeks of age (**Fig 4C**). Analysis of the subjected to loading-to-failure testing (**Fig 4B**) revealed further that *ex vivo* rupture tended to occur more distally and along the inner curvature in WT aortas, but more proximally and ventrally in MFS aortas (**Fig 4D,E**). Moreover, in MFS aortas exposed to BAPN (**Fig 4F**), rupture sites tended to be preferentially located in regions that simultaneously experienced substantial dilatation (mostly >50% increase in local radius) but less substantial thickening (mostly <50% increase in local thickness).

**Figure 4.**
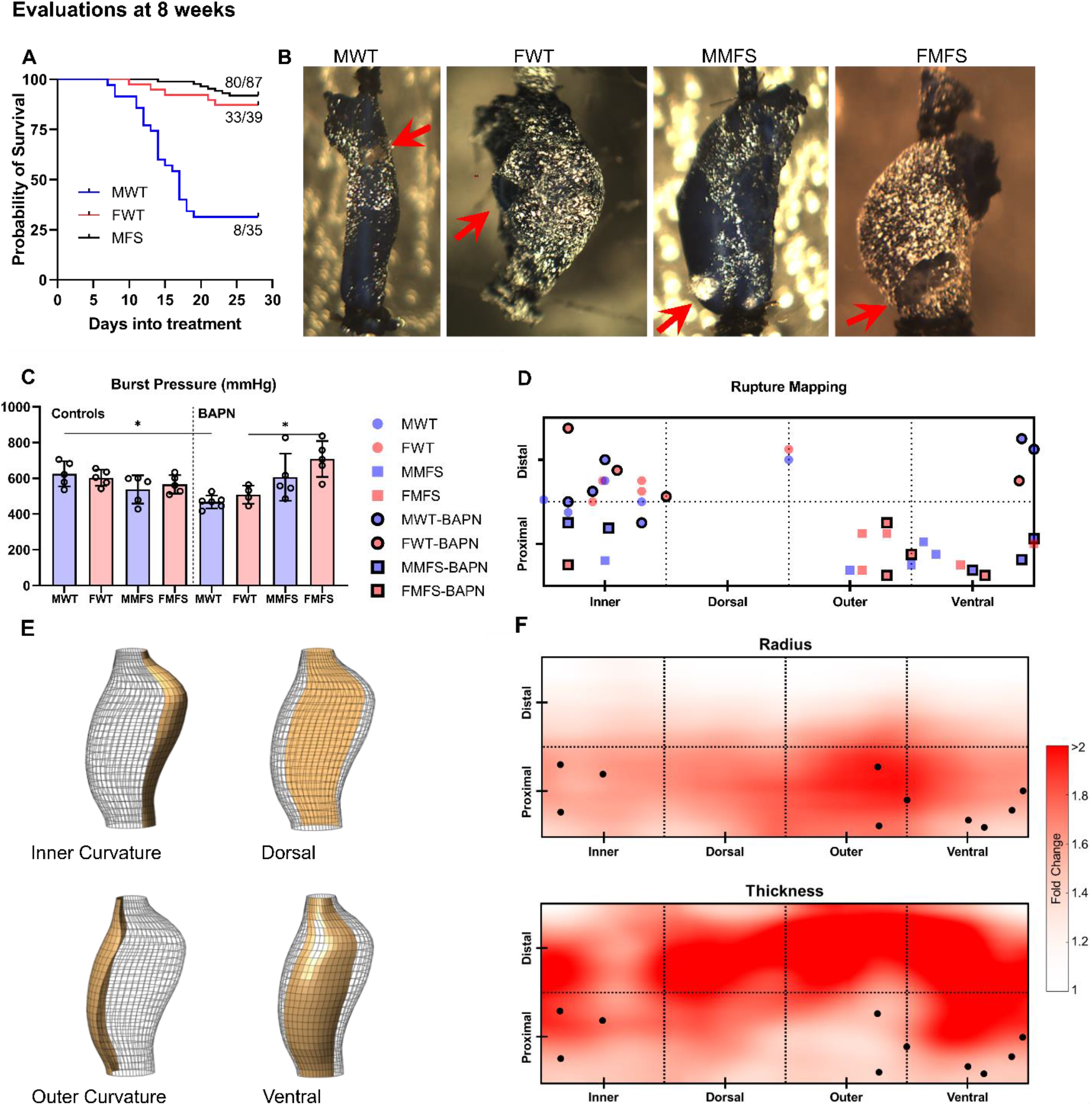
A. Survival curves reveal a dramatically higher BAPN-associated lethality due to aortic rupture in young WT than young MFS mice, particularly in the young male WT mice exposed to BAPN. B,C. Albeit subject to a selection bias (only aortas from surviving mice), *ex vivo* burst-pressure testing confirmed greatest vulnerability in the young male WT aortas (red arrows point to rupture site). D,E. Rupture sites were mapped onto four circumferential (inner curvature, dorsal, outer curvature and ventral) and two axial (proximal half vs. distal half) regions. F. Maps showing the local fold-changes of radius and thickness associated with MFS and BAPN exposure, relative to the WT non-treated controls. Dilatation in the MFS-BAPN group was observed primarily in proximal regions, particularly near the outer curvature, while thickening was mostly observed distally. The locations of failure during *ex vivo* testing of MFS aortas (black points) tended to coincide with regions that experienced local dilatation (i.e., increased radius) *without* substantial thickening.

### Histological and immunohistochemical data reveal key changes in mural collagen following BAPN exposure

Standard histological examinations of fixed aortic cross-sections using picro-sirius red staining and polarized light microscopy further quantified medial and adventitial collagen in all 16 Groups (**Fig 5A-D,F,G**). Among other findings, there was little change in collagen staining in male WT mice following 4-weeks of BAPN that was started at either 4 or 8 weeks of age, but a dramatically higher collagen staining in male MFS mice particularly when BAPN was started at 4 weeks of age. Interestingly, following BAPN exposure, there was also an increase in the ratio of thick:thin adventitial collagen fibers, with a greater increase in thick fibers particularly in the male MFS mice. Importantly, immuno-staining for lysyl oxidase revealed that this important cross-linking enzyme was initially highest in the male MFS aortas and lowest in the male WT mice prior to BAPN exposure, and this difference persisted following BAPN exposure, though at lower levels (**Fig 5A,B,E,H**). Second harmonic generation in non-fixed physiological sections revealed further that adventitial collagen straightness tended not to change with BAPN exposure, but micron-scale collagen fiber bundle widths reduced significantly (**Fig 6A,C,D**). Finally, nano-scale transmission electron micrographs of fixed sections (**Fig 6B**) revealed, among other findings, that the overall area fraction occupied by collagen fibrils in the adventitia decreased dramatically in male WT mice following 4-weeks of BAPN when started at 4 weeks of age but not in the other Groups (**Fig 6E**). Detailed assessments of distributions of fibril cross-sections for all 8 of the 8-week Groups confirm these mean findings while suggesting a pre-existing vulnerability in the male WT aortas (**Fig 6F,G, Fig S11A,B**). In particular, mean fibril cross-sectional area was higher and more variable in MFS aortas compared to their respective WT counterparts; mean fibril area also tended to be larger in female than in male WT mice regardless of BAPN treatment, though this difference was only statistically significant in the BAPN-treated condition due to higher sample sizes. Interestingly, no statistically significant pairwise differences were found in fibril cross-sectional shape as quantified by the aspect ratio, which was remarkably consistent near 1.3±0.2 across all Groups (**Fig S11C, D**). Regardless, the aforementioned trends in macro-scale burst pressures (strength), the lowest of which was in the young male WT vessels (recall **Fig 4C**), correlated well with adventitial collagen fibril cross-sectional area (**Fig 6H**).

**Figure 5.**
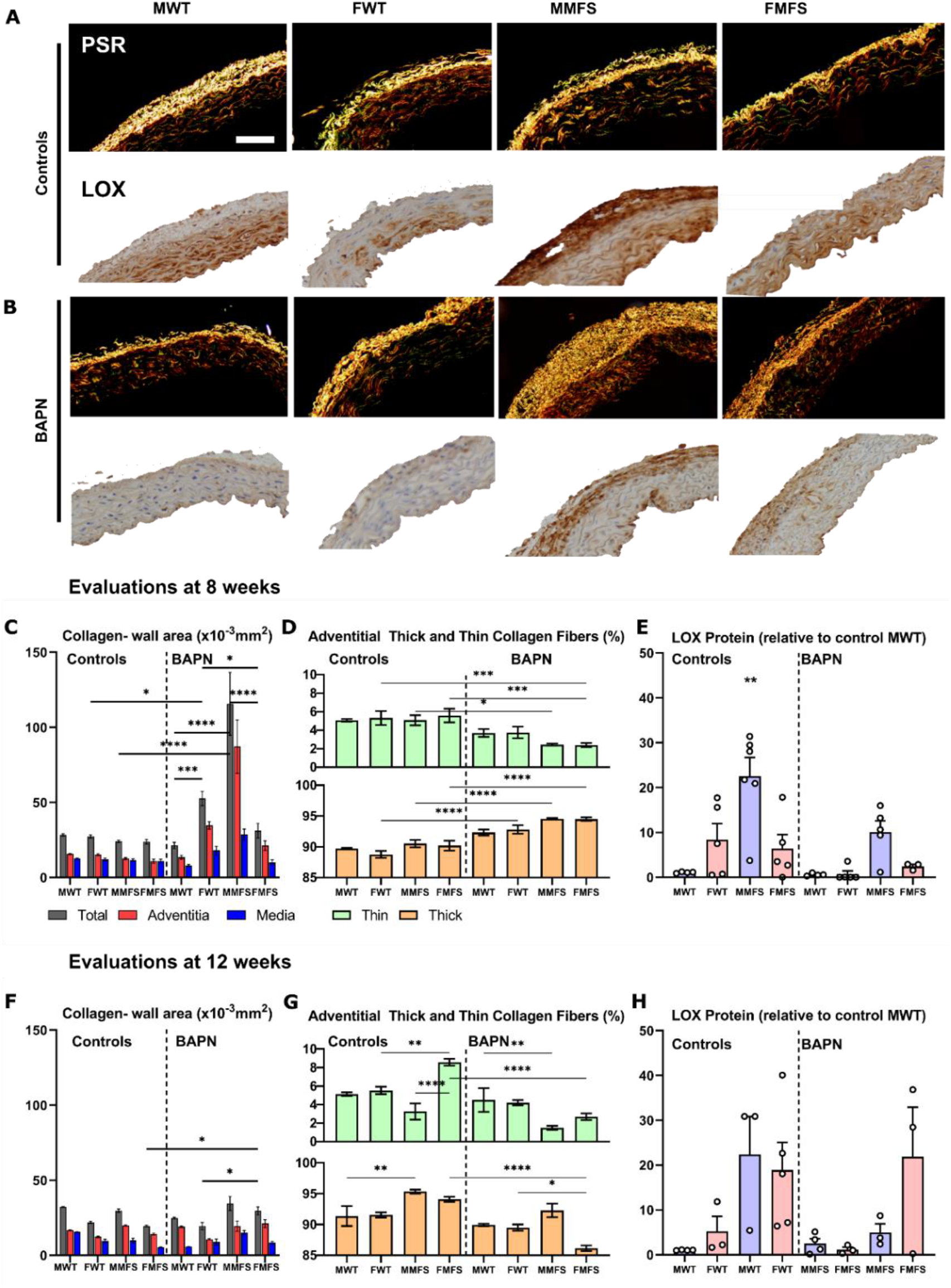
A-D, F,G. Picro-sirius red stained cross-sections revealed modest changes in mural collagen following BAPN exposure in older mice as well as in most younger mice with the stark exception of young male MFS mice in which there was a marked increase in adventitial collagen. Interestingly, BAPN exposure resulted in an increase in the ratio of thick:thin adventitial collagen in all young mice. E, H. Notably, immuno-staining for LOX revealed generally higher levels in female than male mice, and higher levels in MFS than WT mice prior to BAPN exposure. These levels tended to drop following BAPN exposure though group-to-group trends persisted. Statistical comparisons by non-parametric Kruskal Wallis test followed by Dunn’s post-hoc test for multiple comparisons. \*\*\*\**p*<0.0001, \*\*\**p*<0.001, \*\**p*<0.01, \**p*<0.05.

**Figure 6.**
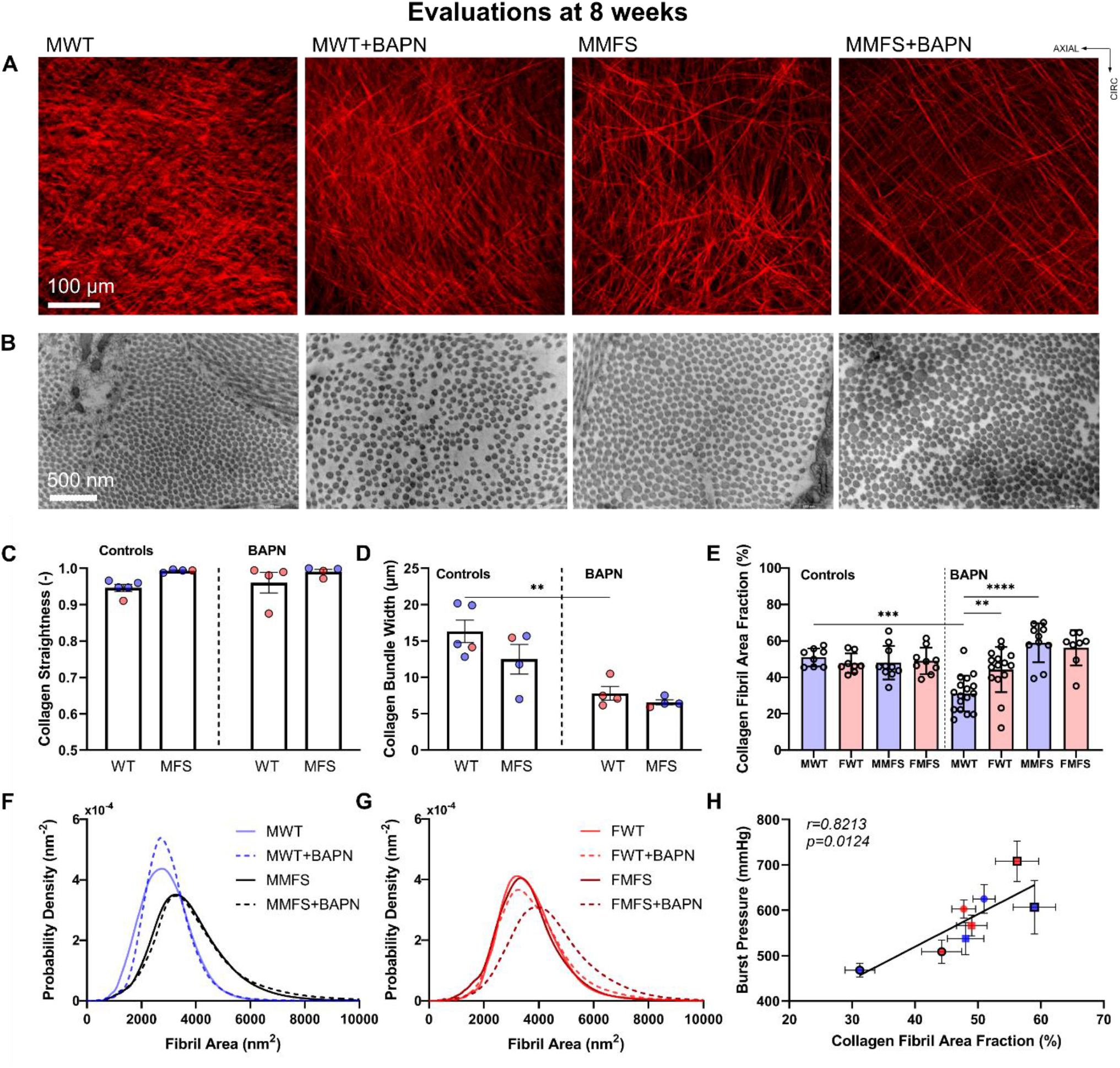
A, C, D. Second harmonic generation in multiphoton microscopy in a sub-set of aortas, from 8-week old mice, revealed reduced collagen bundle diameters (micron scale) following BAPN exposure (significant for the WT) without changes in collagen fiber straightness. E. By contrast, transmission electron microscopy (TEM) revealed in a sub-set of aortas, from 8-week old mice, similar fibril density (nanometer scale) prior to BAPN exposure, independent of sex and genotype, but a marked reduction in fibril density in male WT mice following BAPN exposure. F. In males (both non-exposed and BAPN-exposed), fibril cross-sectional area in MFS mice was higher on average but more variable compared to their respective WT counterparts. G. Mean fibril area in female WT mice tended to be larger than in male WT mice regardless of BAPN exposure, though this difference was only statistically significant in the BAPN-exposed condition due to higher sample sizes (**Fig S11A**). H. Across the 8 groups examined, burst pressure was found to correlate significantly with the collagen fibril area fraction derived from TEM.

## DISCUSSION

It has been known since the 1950s that BAPN can compromise the structural integrity of the aorta, though with a strong dependence on the age at exposure; younger rodents are more vulnerable than older rodents. Most recent studies in mice have given BAPN either to young animals (often starting at weaning, that is, 3 weeks of age) or in combination with induced hypertension in older animals (often 8 weeks of age or older)^31,32^. Given that BAPN only affects newly synthesized fibers, these protocols have proven useful because extracellular matrix turnover is high during aortic development^33^ and hypertension accelerates matrix turnover in the aorta in adults^34^.

The *Fbn1*^*mgR/mgR*^ aorta exhibits a more severe biomechanical phenotype than the *Fbn1*^*C1041G/+*^ aorta^7^, with the former often rupturing spontaneously by 9 weeks of age (soon after biomechanical maturation of the normal aorta) and the latter typically manifesting marked aortopathy only after 9 months^8,9^. There have been attempts to accelerate disease progression in *Fbn1*^*C1041G/+*^ mice, including via chronic exposure to BAPN^28^ and chronic infusion of angiotensin II^35^, noting that infusion of angiotensin II alone leads to thoracic aortic lesions in mature WT and especially *Apoe*^*-/-*^ mice^36,37^. In the prior BAPN study that compared WT and *Fbn1*^*C1041G/+*^ MFS mice (presumably males, though sex was not indicated), ascending aortic diameter increased significantly after 4 and especially 8 weeks of BAPN (daily IP injections) when started in 6-week old MFS, but not in age-matched WT, mice. Our results in 4-week old male WT and MFS mice exposed to BAPN for 4 weeks are generally consistent with these prior findings despite the 2-week difference in the age at which BAPN was started; importantly, however, we also found BAPN-induced dilatation to be absent in young female WT and MFS mice as well as when BAPN was started at 8 weeks of age, independent of sex and genotype. In other words, young male MFS mice were most susceptible to BAPN-exaggerated ascending aortic dilatation with associated reductions in biomechanical functionality.

BAPN was previously given to offset measured increases in *Lox* and *Loxl1* in *Fbn1*^*C1041G/+*^ mice relative to WT controls, with results suggesting that increases in lysyl oxidase activity are compensatory in this MFS model, contributing to its normally slow aneurysmal development^28^. There was no prior assessment of aortic stiffness or strength, however, and limited assessment (light microscopic) of mural composition. Our results on LOX immuno-staining were consistent with this prior study despite the noted differences in study design, with the additional finding herein that LOX trended higher in young female WT compared to age-matched male WT mice and LOX was generally higher in MFS than WT mice independent of age or sex. Although there was no report of increased mortality in the BAPN-treated *Fbn1*^*C1041G/+*^ mice in the prior study, and only low mortality in ours, BAPN significantly increased mortality via aortic rupture when initiated in *Fbn1*^*mgR/mgR*^ mice at 6 weeks of age^28^, consistent with the more severe phenotype of this model.

We initiated BAPN treatment at 4 weeks of age, which is after normal biomechanical maturation of the load-carrying elastic lamellae in WT mice but prior to maturation of intra-lamellar elastic fibers and mural collagen fibers^30,38,39^, to focus first on these possible contributors to wall integrity. Notably, total elastic fiber area inferred via Movat staining, which is dominated by the elastic lamellae, was not reduced in any of the four BAPN-exposed Groups; indeed, it increased slightly when BAPN was started at 4 weeks of age. Although not visible in typical Movat-stained sections, intra-lamellar elastic fibers and fibrillin-1 microfibrils increase progressively in the normal rat aorta from 2+ weeks of age to maturity^38^, which may reflect the emerging need to mechano-sense the increasing wall stress^30^, with mechano-sensing critical to establishing a mechanically homeostatic aortic wall^40,41^. It is also thought that these intra-lamellar elastic fibers / microfibrils may improve the mechanical function or integrity of the lamellae within the media, interacting with other medial matrix components to protect against delamination^15,16^, and to connect the lamellae to the smooth muscle cells to facilitate phenotypic modulation^42–44^. Given the unremarkable gross changes in the elastic lamellae in the young MFS mice following BAPN^39^ plus the recently identified strong correlation between aortic dilatation and elastin porosity in MFS^7^, we used two photon fluorescence to examine elastin porosity in a sub-set of mice. Importantly, we found marked increases in elastin porosity and pore size in young MFS aortas that were exacerbated by BAPN exposure, especially in male MFS, which had the greatest increase in aortic dilatation. Interestingly, these increases in porosity and pore size were greatest in the inner and outer media, the latter often being the site of dissection. Taken together, the present results suggest that compromised intra-lamellar elastic fibers and microfibrils and associated progressive increases in elastin porosity are key drivers of aortic dilatation. A corollary is that, despite little attention in the MFS literature, the intra-lamellar constituents that continue to form throughout development are important contributors to mechanical homeostasis in the normally maturing aorta. The sexually dimorphic BAPN-related dilatation of the MFS aorta, more in young males than females, may suggest that these intra-lamellar elastic fibers and microfibrils develop more slowly in males than in females, thus rendering the male aorta more vulnerable to an age-matched BAPN challenge. These are important observations given that most prior assessments of medial degeneration have focused on breaks in the lamellar elastin^9,17^, primarily in male mice.

Recall that the medial layer (containing nearly all of the elastin and some of the collagen) endows the normal aorta with its compliance (distensibility) and resilience (energy storage capacity) while the adventitia (mostly fibrillar collagen) endows the wall with much of its stiffness and strength, normally serving mainly as a protective sheath that is engaged primarily at elevated hemodynamic loading^45^. We used both biaxial testing at physiological pressures and burst-pressure testing at supra-physiological pressures to assess contributions of these two key layers. Consistent with prior studies, we found that the primary biomechanical characteristics of the *Fbn1*^*C1041G/+*^ MFS aorta were a decreased elastic energy storage capability, which progressed from 8 to 12 weeks of age, and a regionally dependent increased circumferential stiffness that emerged early but did not change dramatically from 8 to 12 weeks^7,8^. Importantly, both changes preceded aneurysmal dilatation. By contrast, circumferential wall stress was not significantly different in WT and MFS aortas under physiological loading at either age, independent of sex. Importantly, in terms of standard geometric and mechanical metrics under physiological loading, BAPN exposure compromised the aorta more in MFS than in WT mice, particularly when was started at 4 versus 8 weeks of age, with the phenotype worse in males than females. Most notable due to early BAPN exposure in MFS mice were the aforementioned (i) marked increases in luminal radius and wall thickness, resulting largely from an accumulation of GAGs and adventitial collagen, and (ii) decrease in elastic energy storage capability, without a noticeable effect on circumferential stiffness. It is not possible to delineate reasons for the further decrease in energy storage, which could include a prevention of intra-lamellar elastic fiber organization^39^ or simply an increase in mural GAGs and collagen that limits distension and thus energy storage. The former is a previously unidentified possibility that deserves further attention. Regardless, the increase in wall thickness helped to reduce biaxial wall stress in the BAPN-challenged MFS aorta, especially when the BAPN was started at 4 weeks of age, independent of sex. Together, the initial similarity in wall stress between WT and MFS aortas combined with the marked decreases in wall stress in the dilating MFS aorta following BAPN exposure focused attention on possible differences in wall strength, and thus collagen.

Remarkably, we found that overall BAPN-related mortality was much greater in young littermate WT (*Fbn1*^*+/+*^) mice than in young MFS (*Fbn1*^*C1041G/+*^) mice, namely 45% vs. 8%, consistent with an early compensatory increase in accumulation and/or cross-linking of collagen in this MFS model^28^. We previously found increased adventitial collagen in another mouse model of thoracic aortic aneurysm and dissection (conditional *Tgfbr1r2* knockout in adults) that appeared to protect against rupture *in vivo* even though this additional collagen differed in structure and function^46^. Most of the WT mice that died of an aortic rupture were young males: 27/35 (77%) males died whereas only 6/39 (15%) females died. Particularly important in this regard, we found that the *ex vivo* burst-pressures (for aortas from surviving animals) were slightly (but not significantly) lower in MFS aortas compared with WT aortas, but this trend reversed following BAPN exposure – burst-pressure was lowest in aortas from BAPN-exposed WT males. Importantly, the regions of failure of the ascending aorta during *ex vivo* testing tended to include the inner curvature and more ventral aspect – distally for WT and proximally for the MFS aortas. Additionally, the regions of *ex vivo* failure in the MFS aorta tended to coincide with regions that experienced dilatation (i.e., increased radius) without substantial thickening, hence supporting the concept that compensatory remodeling is key to protecting the aorta from rupture. Consistent with the findings for burst pressure (strength), recall that immuno-stained LOX was lowest in aortas from WT males, both without and with BAPN exposure. By contrast, LOX was highest in aortas from MFS males, which also showed a marked increase in picro-sirius red detected collagen. There was also an increase in the ratio of thick:thin adventitial collagen fibers following exposure to BAPN, especially in MFS, independent of sex, perhaps suggesting that previously cross-linked thick fibers are more stable. Interestingly, micron-scale reductions in adventitial collagen bundle width were similar in both the rupture-prone young male WT mice and the less-prone young male MFS mice whereas nanometer-scale adventitial collagen fibril density was lower in young male WT than young male MFS aortas (**Fig 6E**), suggesting that it was the fibril level organization that dictated rupture potential (**Fig 6H**).

The strong sexual dimorphism manifested both as greater BAPN-induced dilatation in young male than age-matched female MFS mice and as a much higher pre-mature lethality due to BAPN-induced aortic rupture in young male than age-matched female WT mice. Notably, our choice to start BAPN at P28 is just after the time (~P26) that female mice begin estrus. Although it has been reported that estrogen increases LOX in diverse connective tissues, including bones, skin, and the uterus^47,48^, a recent study using angiotensin II infusion in adult male *Apoe*^*-/-*^ mice suggested that the stronger aortopathy in males was due to the androgens^49^. Importantly, orchiectomy and ovariectomy studies in WT mice suggest that androgens reduce LOX in the male aortas, hence rendering them vulnerable mechanically^50^. Conversely, an elastase + BAPN model of abdominal aortic aneurysms reported greater dilatation in female than male mice, though with low numbers of female mice^51^. There is clearly a need for a detailed study of the remarkable sex differences observed herein and in related models.

Whereas thoracic aortic aneurysms tend to exhibit an increased circumferential material stiffness^52,53^, consistent with disrupted mechano-sensing^11^, it is not clear whether this increase is pathologic or protective. Although BAPN compromised collagen fibers, especially in male WT mice, effects on circumferential stiffness were modest, namely, slight decreases in male and female WT aortas and greater decreases in male and female MFS aortas when BAPN was started at 4 weeks of age with little effect when initiated at 8 weeks of age in the MFS mice. These results suggest that the effect of BAPN on collagen was primarily in reducing strength, not stiffness, perhaps with the former depending more on inter- and intra-molecular cross-links at the fibril level and the latter more on macroscale collagen fiber undulation. Regardless, this critical issue demands further study. Recall, of course, that the exuberant wall thickening in the MFS aortas also reduced stress, with strength > stress critical for aortic integrity. There was also little effect of BAPN on axial material stiffness in the WT aortas though a modest decrease in the MFS aortas, both male and female, when BAPN was initiated at 4 weeks of age. Although the role of axial stiffness is not clear in thoracic aortopathies^52^, the associated reduction in axial wall stress in the MFS aortas in the 4-week BAPN group would again be expected to be protective. We did not assess possible changes in tissue-level contractility, but prior studies have reported decreased myofilaments in aortic smooth muscle cells with BAPN exposure^54^, noting further that the *Fbn1*^*C1041G/+*^ aorta already exhibits decreased contractility^9^. Modest smooth muscle contractility is needed for effective mechano-sensing and mechano-regulation of matrix^11^ and strong vasoconstriction can reduce wall stress and provide some protection for a vulnerable wall^55^, thus there is a need to assess possible reductions in vasoconstrictive capacity in BAPN-exposed aortas, especially given roles of lost smooth muscle contractility in thoracic aortic disease^11,12^.

It is critical to note that inhibiting lysyl oxidase activity via BAPN blocks covalent cross-linking of newly synthesized fibrous matrix rather than breaking cross-links of extant matrix^24^, hence emphasizing the importance of rates of turnover of matrix^56^, which are necessarily higher in developing than mature connective tissue^57^ and higher in hypertensive than normotensive aorta^34^. Increased rates of collagen synthesis can slow aneurysmal enlargement if mechanically competent^58,59^, but can heighten enlargement if not competent^13,28^. Hence, notwithstanding the critical role of compromised elastic fiber integrity in initiating lesion progression in MFS^7^, it is clear that the mechanical functionality of collagen fibers and their turnover are critical modulators of all outcomes – dilatation, dissection, and rupture.

Finally, it has long been known that BAPN can increase intramural GAGs and aggregating proteoglycans^39,60,61^, which have been implicated as key players in dissection and rupture of the thoracic aorta^62–64^. In particular, computational models suggest that, depending on the concentration and distribution of the GAGs, interstitial water sequestered by the GAGs can increase intramural Gibbs-Donnan swelling pressures to levels that can contribute to intramural delamination^65,66^. Consistent with prior studies, we found accumulations of mucoid material in the MFS aortas, which was increased by BAPN exposure. That this increase in GAGs did not increase dissection potential in the MFS aortas reminds us of the key difference between diffuse (as seen herein) and pooled GAGs, with only the latter creating stress concentrations that appear to help nucleate intramural delaminations that lead to dissection.

In summary, taken together, the present results emphasize the high rates of matrix deposition during development and suggest that developing intra-lamellar elastic fibers / microfibrils and mural collagens, especially adventitial, both play critical roles in determining aortic functionality and structural integrity. Although BAPN exacerbated aneurysmal dilatation in the MFS aorta when started during development, it appears that prior compensatory increases in mural collagen and its LOX-mediated cross-linking prevented most of these aortas from progressing to rupture during the 4-week period of study. By contrast, BAPN did not induce aneurysmal dilatation in WT mice, but instead increased rupture-potential especially in the male mice, the aortas of which exhibited the lowest lysyl oxidase activity and the lowest adventitial collagen fibril density. Although thoracic aortic aneurysms and dissections are typically discussed together, the present data clearly delineate different mechanisms of aneurysmal dilatation (related to compromised elastic fibers, including intra-lamellar), here seen mostly in male MFS mice, and transmural rupture (related to compromised collagen fibrils, especially adventitial), here seen mostly in male WT mice with premature mortality resulting from decreases in competent, cross-linked adventitial collagen. Pharmacotherapies directed toward treating MFS patients should seek to preserve collagen integrity while attenuating losses in competent elastic fibers.

## ACKNOWLEGMENTS

This work was supported, in part, by grants from the US National Institutes of Health (P01 HL134605, U01 HL142518). We acknowledge insightful advice from Drs. Dianna M. Milewicz, Martin A. Schwartz, and George Tellides.

## METHODS

### Animals

All live animal care and use conformed to national guidelines and was approved by the Institutional Animal Care and Use Committee (IACUC) of Yale University. Mice, maintained on a C57BL/6J background, were generated by breeding female *Fbn1*^*+/+*^ with male *Fbn1*^*C1041G/+*^ (Jackson Laboratories; stock no. 012885) mice. Littermate *Fbn1*^*+/+*^ (wild-type control, WT) and *Fbn1*^*C1041G/+*^ (Marfan syndrome, MFS) mice were maintained on normal chow and water up to the time of experimental study. To prevent cross-linking of newly synthesized elastic or collagen fibers, female and male WT (FWT, MWT) and female and male MFS (FMFS, MMFS) mice were given β-aminopropionitrile (BAPN; Sigma Aldrich A3134) *ad libitum* at 2000 mg/l in the drinking water for 4 weeks^31^ beginning either at 4 (P28) or 8 (P56) weeks of age. Hence, vessels were excised for testing at either 8 or 12 weeks of age, without or with 4-weeks of BAPN exposure, thus resulting in 16 primary Groups to be compared histo-mechanically. Standard tail-cuff blood pressures were collected prior to euthanasia.

### Biomechanical Testing

We used a series of custom *ex vivo* testing methods to quantify both bulk^67^ and regional^68^ passive mechanical properties of the ascending aorta in the primary material directions, circumferential and axial. Briefly, following euthanasia (CO_2_ inhalation), the ascending aorta was excised from the aortic root to just beyond the brachiocephalic branch. For standard biaxial testing, aortic segments were cannulated with custom glass micro-pipets and placed within a computer-controlled biaxial testing device in a Hank’s buffered salt solution (HBSS) at room temperature to ensure passive responses. The specimens were preconditioned via cyclic pressurization while held at their energetically preferred (*in vivo*) axial stretch, 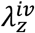, then subjected to a sequence of seven biaxial protocols: cyclic pressurization tests from 10 to 140 mmHg at three fixed axial stretches (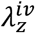 and ±5% of this value) and cyclic axial loading tests from 0 to *f*_*max*_ at four fixed luminal pressures (10, 60, 100, 140 mmHg), where *f*_*max*_ (in mN) was the specimen-specific value reached at a pressure of 140 mmHg and 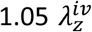. Outer diameter, distending pressure, axial length, and axial force were recorded on-line.

Next, full-field panoramic digital image correlation analyses (pDIC) were used to study regional mechanics. Briefly, the adventitia was stained with Evans blue dye, then gently air-brushed with white India ink to form a unique white speckle pattern over the entire vessel surface (**Fig S12**). The sample, re-cannulated on a co-axial triple-needle assembly, was then placed within a custom 45-degree-angle conical mirror, again submerged within a HBSS-filled bath at room temperature, and connected to a pressure-controlled line. Images containing the reflected speckle pattern on the 45-degree mirrored surface and a static calibration target on the 30-degree outer surface were captured by a nearly vertically-located digital camera (DALSA Falcon 4M30) from eight rotationally symmetric views. Data were collected at 42 quasi-statically loaded configurations: 14 pressures (10-140 mmHg, in 10 mmHg increments) and 3 axial stretches (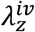 and ±5% of this value, as in standard biaxial testing) for a total 336 images per specimen. The reference configuration was defined at a pressure of 80 mmHg and *in vivo* axial stretch. Custom MATLAB scripts were used to unwrap the pDIC images such that the polar angle and radial coordinates in the acquired image were mapped onto a rectangular grid. Serial correlations based on a normalized cross-correlation approach were then performed between pairs of unwrapped images over all deformed configurations. Combining a direct linear transformation (DLT)-based calibration for each image with knowledge of the conical mirror geometry, a ray-tracing procedure was used to reconstruct the physical location of each correlated point over the sample surface in the coordinate system of the conical mirror. In the end, this procedure allowed reconstruction and meshing of the 3D surface geometry at each deformed configuration and thereby allowed straightforward computation of full-field surface deformations. Local OCT-derived thickness measurements collected at 100 cross-sections along the length of the vessel were also mapped onto the pDIC reconstructed geometry using an automatic co-registration pipeline described previously^69^. Detailed information about our multimodality pDIC and OCT pipeline can be found in our prior papers^68,70,71^. Finally, in a sub-set of mice, the vessels were then placed within another experimental system and pressurized at their *in vivo* axial length until failure.

### Material Characterization

The passive nonlinear mechanical properties were quantified using an independently validated^72^ constitutive relation that we have shown describes well the biaxial mechanical behavior of the mouse aorta^52,73^. First, the unloading portion of the last cycle of all seven pressure-diameter and axial force-length tests were fit simultaneously with a four-fiber family constitutive model using a Levenberg-Marquardt nonlinear regression of the data. Using data during unloading reveals the elastic energy stored during deformation that would be available to work on the distending fluid. This four-fiber family model is written in terms of a scalar stored energy function *W*, namely

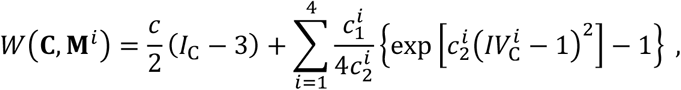

where the eight model parameters to be determined via regression are 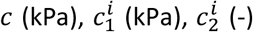, and 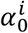 (radians), with *i* = 1,2,3,4 denoting the four predominant fiber family directions: axial 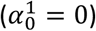, circumferential 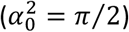, and symmetric diagonal 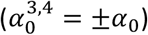. *I*_*C*_ = *tr*(**C**) and 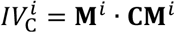 are coordinate invariant measures of the finite deformation, with the right Cauchy-Green tensor **C** = **F**^T^**F** computed from the experimentally measured deformation gradient tensor **F** = diag[*λ*_*r*,_ *λ*_*θ*,_ *λ*_*z*_], with *det***F** = 1 because of assumed incompressibility. The direction of the *i*^*th*^ family of fibers was prescribed by 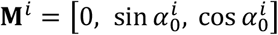, with 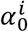 denoting a fiber angle relative to the axial direction in the traction-free reference configuration. Values of mean biaxial wall stress and material stiffness were computed from the stored energy function and calculated at a common pressure of 100 mmHg.

Second, the pDIC data were used to estimate best-fit parameters of our four-fiber constitutive relation locally, herein at ~1000 sites around the circumference and along the length of each specimen: 40 circumferential patches defined by Θ_m_, *m* ∈ [1,40], × 25 axial patches defined by *Z*_n_, *n* ∈ [1,25], with a cylindrical coordinate system defined over Θ ∈ [−π, π] and Z ∈ [0, L], where L is the length of the sample in the *in vivo* relevant reference configuration. Using the pDIC-derived local displacement fields, Green strains (**E** = (**C −I**)/2, where **C** is the right Cauchy-Green tensor and **I** is the second-order identity tensor) were calculated at each Gauss point of a 4-noded rectangular surface patch assuming (for simplicity) an incompressible neo-Hookean strain energy function. Strains were calculated using the open-source finite element software, FEBio. Then, the material behavior at each Gauss point was modeled with the aforementioned four-fiber family *W*. The principle of virtual power was enforced at each Gauss point to achieve inverse characterization within each Θ_m_*Z*_n_ element. The unknown material parameters were iteratively updated for each element using a derivative-free genetic algorithm followed by a gradient-based local optimization to maximize the log-likelihood of observing the experimentally measured pressures and axial loads^69^. Once completed, the final set of the identified parameters at each element was used to compute full-field distributions of the different mechanical metrics, as described previously. For additional details, see our prior papers^53,69,71^.

To perform spatially local statistical analyses of group-wise differences in geometric and mechanical properties, we developed an automatic pipeline parametrize and co-register all pDIC-derived geometries in our database, extending similar approaches for other tissues with genus-1 topology^74^. Starting from each vessel-specific quadrilateral patch surface obtained via pDIC-based reconstruction, we performed a principal component analysis of the nodal coordinates and automatically identified two objective landmarks on the vessel (**Fig S13**): (1) the outermost point on the outer curvature and (2) the ligation point on the distal arch. To control for differences in vessel length, we linearly transformed the length scale along the axial direction such that a corresponding size-independent “axial coordinate” had a value of 0 at the outermost point on the outer curvature and a value of 1 at the arch ligation point (**Fig S13**). Because the vessel geometry goes past these locations, the resulting axial coordinate values extend beyond the [0, 1] interval. One sample (third male WT+BAPN specimen shown in Fig 3B) was excluded from this analysis alone since severe dilation precluded identification of the required anatomic landmarks. We then also parametrized the vessel circumferentially on the interval [−1, 1] using the in-plane angle of each patch node with respect to the local centerline along the vessel, with periodicity at −1 and 1 (which corresponds to the middle of the outer curvature). For co-registration and analysis, we re-discretized the surface within the “correspondent” region along the axial direction that was shared by all vessels included in the analysis. Within this shared region, to facilitate interpretation of all results presented, the vessel wall surface was segmented into anatomically relevant regions, namely “proximal” and “distal” along the axial direction, and “inner,” “dorsal,” “outer,” and “ventral” around the circumference (**Fig S13**).

At each mesh point within the correspondent region of the vessel wall, we quantified the effects of sex, genotype, and BAPN exposure on the local radius and thickness of the vessel. Denoting the metric of interest (radius or thickness) as *y*, we modeled the group- and location-specific log-values as a linear function of the predictors using

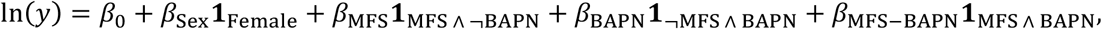

where **1** is the indicator function for the subscripted condition(s), “∧” denotes the logic AND function, and “¬” denotes the logic NOT function. The best-fit parameters *β*_*i*_ are estimated via least-squares regression of ln(*y*). Exponentiating both sides, the relation becomes

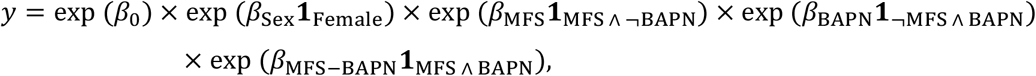

where the “baseline” exp (*β*_0_) corresponds to the predicted value for MWT non-treated mice, exp (*β*_Sex_) is the fold-change when the sex is changed to female (within any strain/treatment group), exp (*β*_MFS_) is the fold-change for MFS without BAPN treatment, exp (*β*_BAPN_) is the fold-change for BAPN treatment without MFS, and exp (*β*_MFS−BAPN_) is the fold-change for MFS coincident with BAPN treatment.

### Multiphoton microscopy

In a subset of 8-week old mice, following standard mechanical testing, vessels were immersed in HBSS at room temperature, and held at a diastolic configuration, i.e., specimen specific axial stretch and a common intraluminal pressure of 80 mmHg. A LaVision Biotec TriMScope two-photon microscope operated with a TiSa laser tuned at 820 nm was used and outfitted with an Olympus 20X objective lens (N.A. 0.95) oriented transversely to the surface of the aortic sample. Three signals were: second harmonic generation (SHG) for fibrillar collagen (390-425 nm), two-photon excited fluorescence for elastin (500-550 nm), and SYTO17 red fluorescence of cell nuclei (above 550 nm). Three-dimensional images were acquired with an axial-circumferential field of view of 500 μm × 500 μm at a consistent anatomical location corresponding to the center of the outer curvature, numerical imaging resolution of 0.48 μm/pixel, and out-of-plane (radial axis) step size of 1 μm/pixel. All images were processed as previously described^7^. Briefly, layer-specific cell densities were calculated as the number of cells per unit volume by counting the number of nuclei within defined sub-volumes of the 3D image and normalizing by the appropriate layer-specific thickness. Three cell groups were considered based on characteristic shapes of their nuclei and radial location: endothelial (monolayer on intima), smooth muscle (within the media), and cells within the adventitia. The three-dimensional elastin porosity was defined as the ratio between the volume identified as voids and the volume occupied by elastic fibers – including lamellar and intra-lamellar. For this purpose, an adaptive 3D filter was applied to the remaining volume to reduce possible inhomogeneities due to local waviness of the elastic lamellae, the image was binarized using the automatic local Phansalkar thresholding method, and the volume of the identified voids was quantified as a black-to-white ratio. The description of the collagen fiber bundles focused on in-plane (i.e., axial-circumferential plane of the artery) parameters of fiber straightness and fiber bundle width. Straightness was computed as the ratio of end-to-end to total fiber length for fibers consistently selected at multiple positions. Bundle width was measured as the transversal section of multiple bundles of fibers. Additional details can be found in our previous paper^7^.

### Histology and Transmission Electron Microscopy (TEM) - Sample preparation

Following the multi-step mechanical testing protocol, vessels were prepared for either standard immuno-histological examination or transmission electron microscopy (TEM). In the former, segments were fixed overnight in a 10% neutral buffered formalin solution and stored in 70% ethanol at 4°C until histological processing. Samples were embedded in paraffin and serially cut across the vessel cross-section at a thickness of 5 µm. Mounted slides were then stained with: 1) Movat’s pentachrome (MOV) to detect elastin in black, collagen in yellow/brown, cytoplasm in purple, aggregating glycosaminoglycans/proteoglycans (GAGs/PGs) in aqua/blue, and fibrin in pink/red and 2) picro-sirius red (PSR) to detect fibrillar collagens in a spectrum from red-to-green (**Fig S14**). Additional sections were immuno-stained with primary antibody against LOX (ab174316 Abcam) to detect (in brown) lysyl oxidase. High-resolution images were acquired using an Olympus BX/51 microscope at 20X magnification. Note that MOV images were collected under standard bright-field conditions while PSR images were collected as dark-field images using polarized light to detect variations in collagen fiber birefringence, with separate exposure times to capture medial and adventitial collagens. MOV sections were used to compute medial and adventitial wall areas, as well as wall areas for elastin, cytoplasm, collagen, GAGs/PGs, and fibrin. Medial and adventitial collagen wall areas were quantified separately from PSR sections at their respective exposures using pixel-based analysis; collagen fibers in the total wall were then categorized as thick (red/orange) or thin (yellow-green) in medial exposure images using color-based analysis routines. Area fractions were computed on a layer-specific basis using wall areas of elastin, cytoplasm, GAGs/PGs, and fibrin from MOV and wall areas of collagen from PSR. For TEM, aortic segments were fixed in 2.5% glutaraldehyde/2% paraformaldehyde in sodium cacodylate buffer at room temperature for 2 h at 4°C. Samples were then rinsed in sodium cadodylate buffer before post-fixation in 1% osmium tetraoxide for 1 h and subsequent staining using 2% uranyl acetate for 1 h. Samples were washed and embedded in resin and viewed using a FEI Technai Biotwin electron microscope^46^.

### TEM – analysis

To analyze the spatial distribution, size, and shape of collagen fibrils within each experimental group, we developed a novel semi-automated pipeline to segment individual fibrils from each transmission electron micrograph (83 images total, with several hundred fibrils in each). As an initial preprocessing step, the user interactively selects regions of the image to exclude from the analysis—for example, areas containing cellular material or GAGs, rather than collagen fibrils (**Fig S15A,B**). The image is then smoothed using a Gaussian filter with bandwidth equal to 0.3% of the smallest dimension (height or width) of the image (**Fig S15B**). To binarize the filtered image into fibril and non-fibril pixels, a bivariate quadratic function *f*_1_(*u, v*) is first fit to the intensity values *I*_filt_(*u, v*) using robust least-squares regression with a Cauchy weight function, where (*u, v*) are pixel coordinates. The modeled intensity values *f*_1_(*u, v*) are adopted as a local adjustment factor, and the intensity values of the filtered image are adjusted using

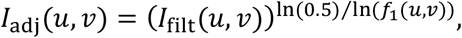

such that pixels with intensity *I*_filt_(*u, v*) = *f*_1_(*u, v*) are mapped to 0.5. This spatially heterogeneous (yet smooth) adjustment attenuates the influence of regional variations in image intensity (e.g., due to non-uniform lighting) during the binarization step. Using Otsu’s method^75^, the transformed image is then binarized (*I*_bin_) according to an optimal threshold (**Fig S15C**), with dark pixels categorized as fibrils.

For each pixel in *I*_bin_, the Euclidean distance to the nearest non-fibrillar pixel is computed, producing a distance field *D* with the same dimensions as *I*_bin_. Then, for different Gaussian filter bandwidths *σ*_*i*_ (ranging from 0.5 to 10 pixels, in steps of 0.5 pixels), the number of local peaks *N*(*σ*_*i*_) in *D*_filt_(*σ*_*i*_)—taken to be the number of “detected” fibrils—is computed. A 5th-degree polynomial *p*(*σ*) is fit to the resulting (*σ*_*i*_, *N*(*σ*_*i*_)) data, from which the optimal bandwidth *σ*^∗^ is considered to be the lowest positive *σ* where (*dp*/*dσ*)^2^ reaches a local minimum. Although *p*(*σ*) is generally monotonically decreasing within the interval of interest, *σ*^∗^ corresponds to the lowest positive *σ* where the slope of *p*(*σ*) is locally the nearest to 0, meaning that the number of detected fibrils is minimally sensitive to the filter bandwidth at this point. The first estimate of the fibrils’ centroid coordinates are thus the locations of the local peaks in *D*_filt_(*σ*^∗^), with the corresponding Voronoi cells defining a first estimate of the fibril “neighborhoods” (**Fig S15D**).

At this stage, the user may interactively add, remove, or modify fibril centroids that were undetected, erroneously detected, or poorly estimated, respectively, if needed. For each Voronoi cell, if any of the cell’s vertices are either outside of the image area or within a previously excluded region, the corresponding fibril is labeled as a “boundary fibril,” to be excluded from future analysis results (e.g., cross-sectional area computations) since part of the fibril is mostly likely not visible.

To improve upon the initial image binarization, a “characteristic intensity” is defined for each Voronoi cell, equal to the mean intensity of pixels in that cell whose pixel-to-centroid distance is less than the 5th percentile of pixel-to-centroid distances within that cell. A smooth interpolant *f*_2_(*u, v*) of the characteristic intensities (defined at the centroids for the purposes of interpolation) is then computed, after which the adjusted image is *readjusted* using

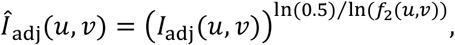

and *Î*_adj_ is binarized (again using Otsu’s method) to yield *Î*_bin_. To summarize the overall presence of collagen fibrils, we compute the fibril area fraction as the percentage of *I*¬bin consisting of fibril pixels.

The newly segmented fibril pixels are then clustered by fitting a 2D Gaussian mixture model to the fibril pixel coordinates, using the current centroid locations as an initial guess for the *N* cluster means and the Voronoi cell memberships as initial cluster assignments. After fitting the model, posterior membership probability fields *P*_*i*_(*u, v*) are computed over the entire image for each cluster (i.e., each fibril) *i*, and the fibril neighborhood boundaries are redefined as *P*_*i*_(*u, v*) = 0.5 (**Fig S15E**). The cross-sectional area of each fibril is computed by summing the areas of all fibrillar pixels within the corresponding neighborhood boundary. At the group level, distributions of fibril area are computed via kernel density estimation of log-transformed area values to enforce a strictly positive support. Under the approximation that fibril pixel coordinates belonging to a particular fibril are uniformly distributed within an ellipse, the major and minor radii can be computed as 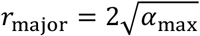 and 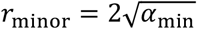 respectively^76^, where *α*_max_ and *α*_min_ are the maximum and minimum eigenvalues of the covariance matrix for those pixel coordinates (**Fig S15F**). Likewise, the orientation of the ellipse is taken from the eigenvectors of the covariance matrix. As a final correction step, the user may interactively remove any ellipses that fit the actual fibril boundary poorly.

### Statistical Methods

Due to deviations from normality (data failed to pass Kolmogorov-Smirnov normality test), we used the non-parametric Kruskal Wallis one-way ANOVA on Ranks followed by Dunn’s post-hoc test for multiple comparisons to compare results across study Groups. Correlations, where appropriate, were assessed using the non-parametric Spearman’s rank correlation coefficient, *r*. The Kaplan-Meier product limit estimator was used to analyze survival distributions. For all reported comparisons, a value of *p* < 0.05 was considered significant. Multiple levels of statistically significant differences across Groups are indicated in each figure, as appropriate.

