## Supplementary Material for "Compensatory aortic remodeling in Marfan syndrome protects against sexually dimorphic rupture during a BAPN challenge"

[2]Vascular Biology and Therapeutics Program
Yale School of Medicine, New Haven, CT

Running title: Age, Sex, and ECM Cross-Linking in MFS

Address for Correspondence:

J.D. Humphrey, Ph.D.
Department of Biomedical Engineering
Yale University, New Haven, CT 06520 USA

+1-203-432-6528

### SUPPLEMENTAL TABLES

**Table S1.** Key geometric and mechanical metrics from all 8 groups evaluated at 8 weeks of age, including those mice without and with preceding 4-week exposures to BAPN. The pressure-dependent quantities were evaluated at 100 mmHg given the lack of a measured difference in tail-cuff blood pressures.

| | **Evaluations at 8 weeks** | | | | | | | |
|---|---|---|---|---|---|---|---|---|
| | **No treatment** | | | | **BAPN-exposed** | | | |
| | **MWT** | **FWT** | **MMFS** | **FMFS** | **MWT** | **FWT** | **MMFS** | **FMFS** |
| | **n =5** | **n = 5** | **n = 5** | **n = 5** | **n = 5** | **n = 5** | **n = 5** | **n = 5** |
| **Unloaded dimensions** | | | | | | | | |
| Wall Thickness (μm) | 125 ± 8 | 141 ± 13 | 123 ± 7 | 129 ± 8 | 129 ± 6 | 125 ± 5 | 257 ± 43 | 189 ± 23 |
| Outer Diameter (μm) | 1103 ± 29 | 1131 ± 9 | 1216 ± 33 | 1173 ± 21 | 945 ± 35 | 1013 ± 29 | 1784 ± 268 | 1289 ± 134 |
| **Loaded dimensions** | **P=100mmHg** | | | | | | | |
| Outer Diameter (μm) | 1738 ± 42 | 1760 ± 36 | 1908 ± 64 | 1788 ± 33 | 1553 ± 37 | 1532 ± 49 | 2684 ± 288 | 1907 ± 160 |
| Wall Thickness (μm) | 42 ± 3 | 49 ± 5 | 41 ± 2 | 45 ± 3 | 42 ± 4 | 48 ± 3 | 123 ± 29 | 85 ± 14 |
| Inner Radius (μm) | 827 ± 22 | 831 ± 19 | 913 ± 32 | 849 ± 16 | 735 ± 19 | 718 ± 25 | 1219 ± 144 | 869 ± 80 |
| *in vivo* Axial Stretch ($\lambda_z^{iv}$) | 1.72 ± 0.04 | 1.67 ± 0.04 | 1.63 ± 0.04 | 1.69 ± 0.03 | 1.68 ± 0.04 | 1.57 ± 0.06 | 1.32 ± 0.10 | 1.38 ± 0.09 |
| *in vivo* Circumferential Stretch ($\lambda_\vartheta$) | 1.73 ± 0.02 | 1.73 ± 0.04 | 1.69 ± 0.03 | 1.67 ± 0.01 | 1.86 ± 0.06 | 1.67 ± 0.04 | 1.73 ± 0.08 | 1.68 ± 0.07 |
| **Cauchy Stresses (kPa)** | | | | | | | | |
| Circumferential, $\sigma_\vartheta$ | 267 ± 16 | 238 ± 31 | 283 ± 17 | 253 ± 14 | 229 ± 16 | 204 ± 17 | 156 ± 26 | 150 ± 22 |
| Axial, $\sigma_z$ | 236 ± 18 | 203 ± 25 | 235 ± 12 | 208 ± 11 | 216 ± 18 | 167 ± 17 | 88 ± 24 | 107 ± 22 |
| **Linearized Stiffness (MPa)** | | | | | | | | |
| Circumferential, $C_{qqqq}$ | 1.56 ± 0.11 | 1.45 ± 0.22 | 3.04 ± 0.28 | 2.53 ± 0.16 | 1.43 ± 0.18 | 1.41 ± 0.15 | 2.27 ± 0.59 | 2.98 ± 0.39 |
| Axial, $C_{zzzz}$ | 1.10 ± 0.07 | 0.94 ± 0.11 | 1.33 ± 0.07 | 1.15 ± 0.07 | 1.20 ± 0.08 | 0.93 ± 0.07 | 0.47 ± 0.11 | 0.83 ± 0.10 |
| **Stored Energy (kPa)** | 82 ± 3 | 72 ± 6 | 63 ± 4 | 55 ± 3 | 62 ± 4 | 49 ± 6 | 22 ± 6 | 21 ± 5 |

**Table S2.** Similar to Table S1 except for the 8 groups of mice evaluated at 12 weeks of age.

| | **Evaluations at 12 weeks** | | | | | | | |
|---|---|---|---|---|---|---|---|---|
| | **No treatment** | | | | **BAPN-exposed** | | | |
| | **MWT** | **FWT** | **MMFS** | **FMFS** | **MWT** | **FWT** | **MMFS** | **FMFS** |
| | **n = 5** | **n = 5** | **n = 5** | **n = 5** | **n = 5** | **n = 5** | **n = 5** | **n = 5** |
| **Unloaded dimensions** | | | | | | | | |
| Wall Thickness (μm) | 135 ± 6 | 132 ± 6 | 144 ± 18 | 140 ± 7 | 129 ± 2 | 113 ± 3 | 183 ± 20 | 164 ± 15 |
| Outer Diameter (μm) | 1098 ± 88 | 1144 ± 24 | 1320 ± 146 | 1094 ± 42 | 1022 ± 40 | 1022 ± 38 | 1470 ± 174 | 1215 ± 113 |
| **Loaded dimensions** | **P=100mmHg** | | | | | | | |
| Outer Diameter (μm) | 1756 ± 24 | 1718 ± 16 | 1918 ± 141 | 1740 ± 28 | 1679 ± 42 | 1625 ± 93 | 2130 ± 168 | 1946 ± 195 |
| Wall Thickness (μm) | 48 ± 2 | 47 ± 4 | 58 ± 2 | 49 ± 3 | 41 ± 1 | 40 ± 2 | 82 ± 17 | 63 ± 8 |
| Inner Radius (μm) | 830 ± 22 | 812 ± 19 | 901 ± 32 | 821 ± 16 | 799 ± 19 | 773 ± 25 | 983 ± 144 | 910 ± 80 |
| *in vivo* Axial Stretch ($\lambda_z^{iv}$) | 1.77 ± 0.03 | 1.70 ± 0.04 | 1.56 ± 0.05 | 1.64 ± 0.02 | 1.71 ± 0.04 | 1.64 ± 0.06 | 1.48 ± 0.08 | 1.48 ± 0.07 |
| *in vivo* Circumferential Stretch ($\lambda_\vartheta$) | 1.69 ± 0.02 | 1.67 ± 0.01 | 1.60 ± 0.10 | 1.74 ± 0.05 | 1.84 ± 0.05 | 1.73 ± 0.04 | 1.63 ± 0.08 | 1.79 ± 0.03 |
| **Cauchy Stresses (kPa)** | | | | | | | | |
| Circumferential, $\sigma_\vartheta$ | 205 ± 34 | 240 ± 19 | 209 ± 23 | 215 ± 17 | 262 ± 5 | 263 ± 28 | 179 ± 22 | 196 ± 9 |
| Axial, $\sigma_z$ | 204 ± 2 | 220 ± 16 | 156 ± 15 | 185 ± 12 | 244 ± 12 | 234 ± 20 | 128 ± 23 | 156 ± 19 |
| **Linearized Stiffness (MPa)** | | | | | | | | |
| Circumferential, $C_{qqqq}$ | 1.38 ± 0.06 | 1.50 ± 0.12 | 2.84 ± 0.62 | 2.13 ± 0.09 | 1.58 ± 0.08 | 1.62 ± 0.19 | 3.11 ± 0.43 | 2.37 ± 0.26 |
| Axial, $C_{zzzz}$ | 1.07 ± 0.02 | 1.06 ± 0.07 | 1.05 ± 0.21 | 1.05 ± 0.05 | 1.22 ± 0.06 | 1.25 ± 0.06 | 0.86 ± 0.14 | 0.98 ± 0.09 |
| **Stored Energy (kPa)** | 71 ± 7 | 70 ± 3 | 37 ± 1 | 48 ± 4 | 74 ± 3 | 65 ± 6 | 29 ± 7 | 35 ± 5 |

**SUPPLEMENTAL FIGURES of RESULTS**

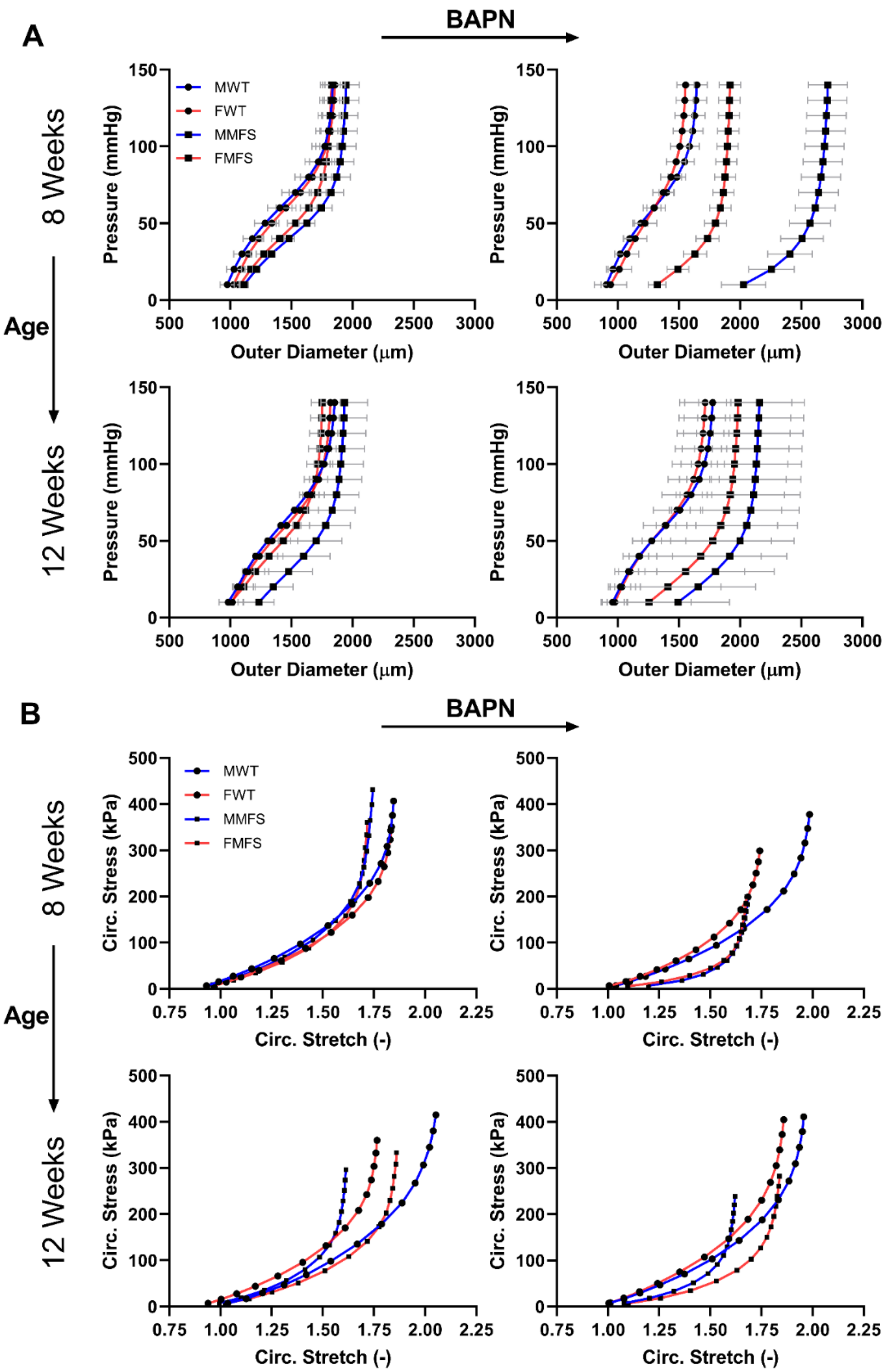


**Figure S1.** A. Pressure-diameter relations for all 16 groups: female (F) and male (M), wild-type (WT) and *Fbn1*$^{C1041G/+}$ Marfan (MFS) ascending aortas with BAPN given for 4 weeks beginning either at 4 weeks of age (then evaluated at 8 weeks) or at 8 weeks of age (then evaluated at 12 weeks of age), with age- and sex-matched controls not receiving BAPN. B. Related circumferential Cauchy stress-stretch relations for the same 16 groups. Note the emergence of a MFS phenotype, characterized by aortic dilatation.

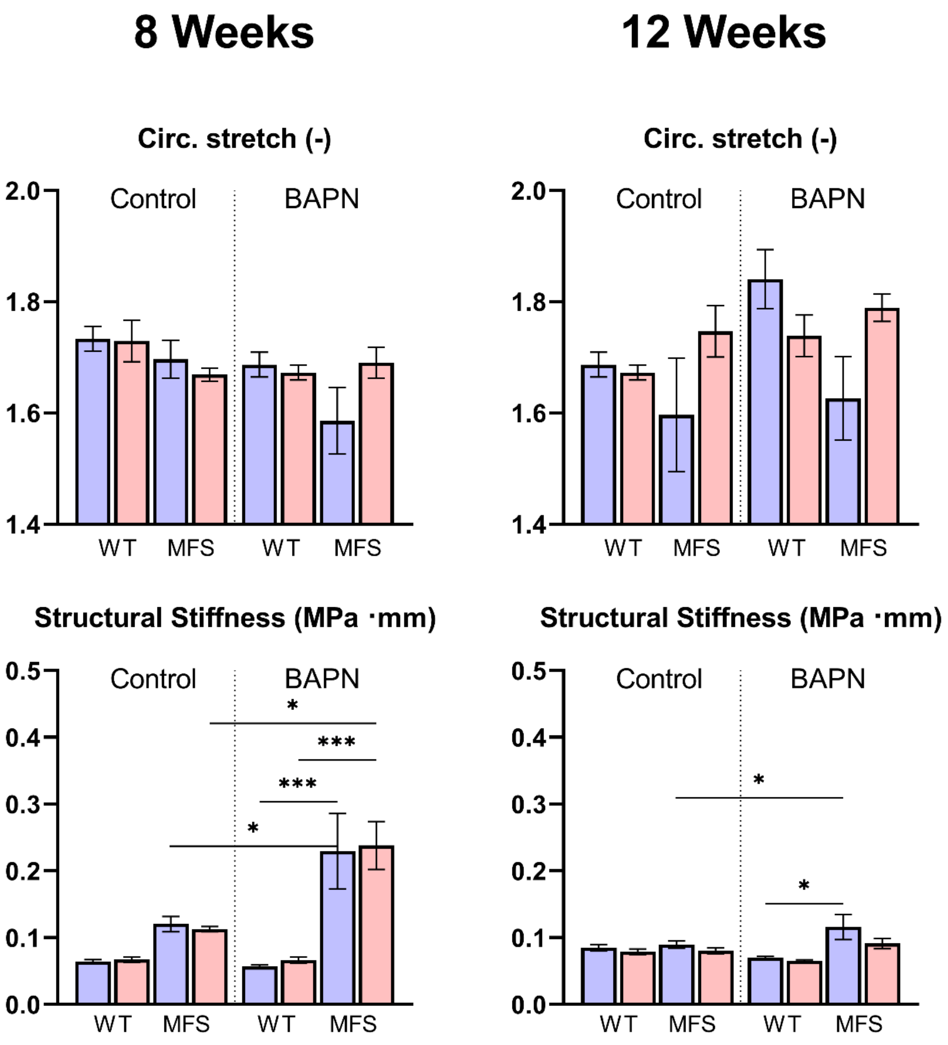


**Figure S2**. Additional (see Fig 1 in the main text) mechanical metrics inferred from the *ex vivo* biaxial testing of all 16 groups. Note the significant increase in the structural stiffness of ascending aorta from the young BAPN-exposed MFS mice (Bottom-left), which corresponds to a dramatic decrease in the arterial distensibility (Top-left).

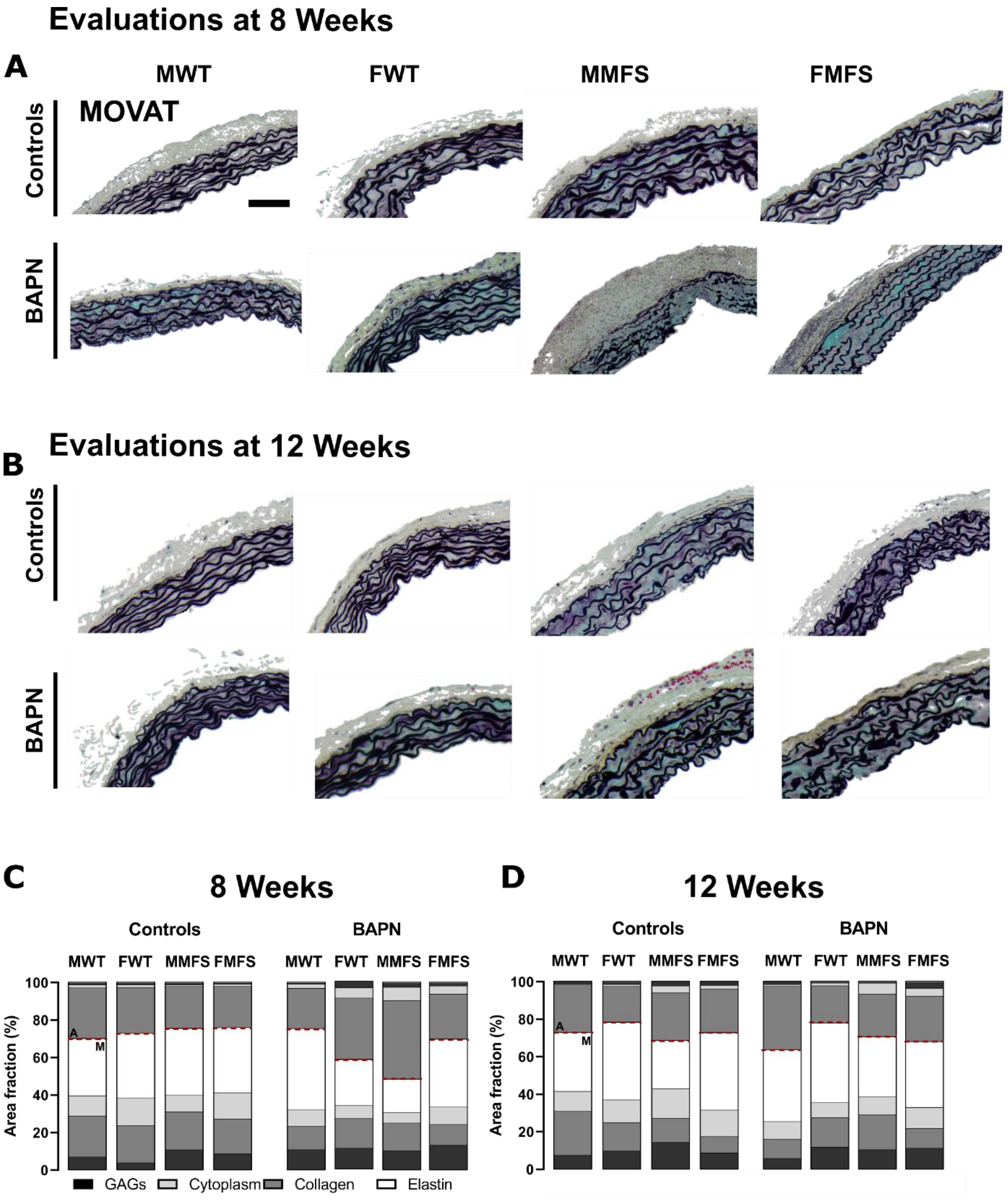


**Figure S3.** A, B. Representative portions of Movat-stained histological cross-sections from all 16 groups. Elastic fibers are shown in black, glycosaminoglycans in blue/aqua, collagen in yellow/brown. C,D. Quantification of mural percentages separately by medial (M) and adventitial (A) layers. Note, in particular, the marked increase in adventitial fraction in the young male MFS aortas following BAPN exposure.

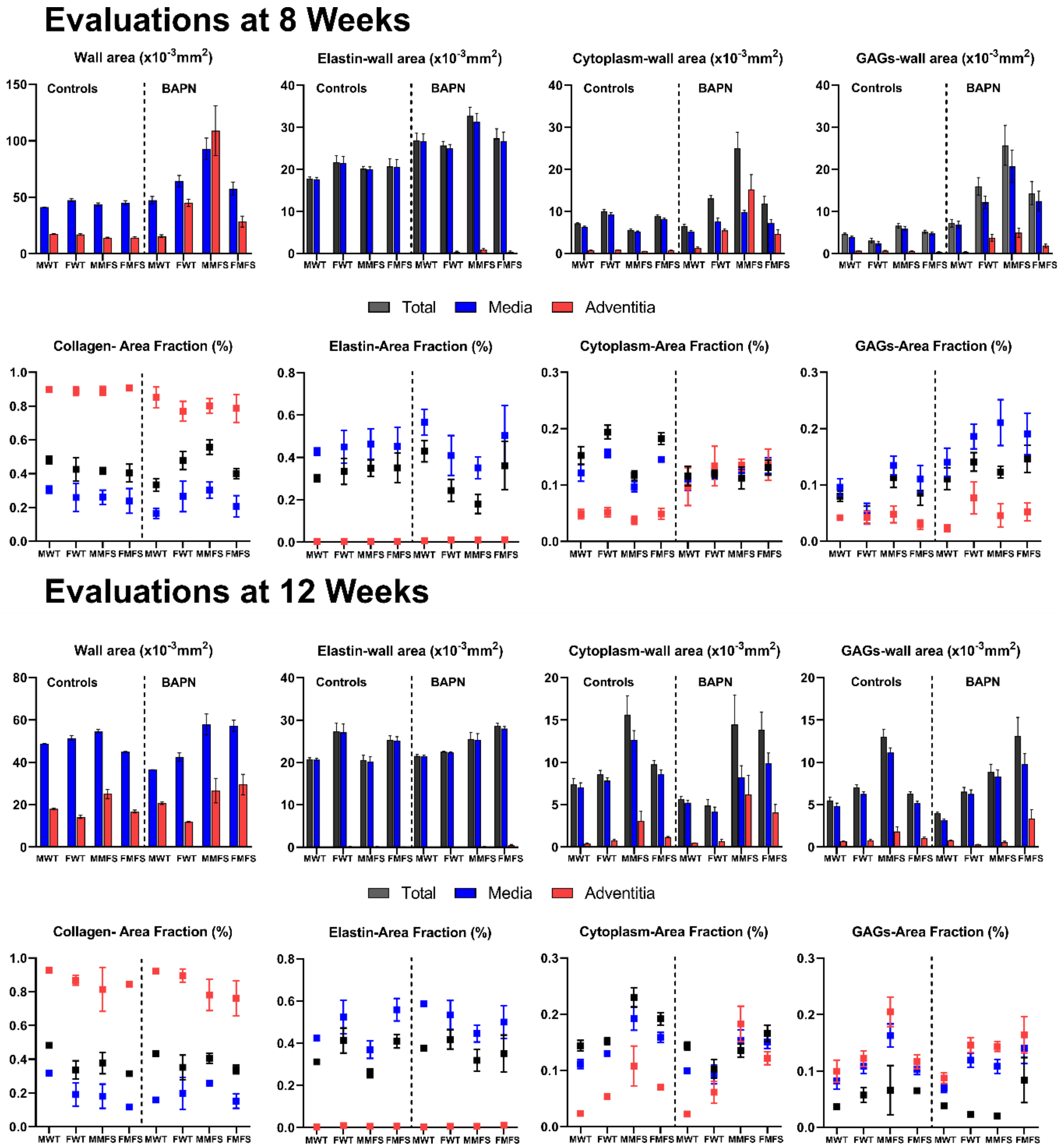


**Figure S4**. A companion to Fig S3, shown here is additional layer-specific quantification of data from Movat-stained sections for all 16 groups. Data are presented for wall area and area fraction with mean ± standard deviation. Note the increase of adventitial cytoplasm in the BAPN-exposed, 8-week old groups, particularly the male MFS, consistent with the multiphoton observations (Fig 2A).

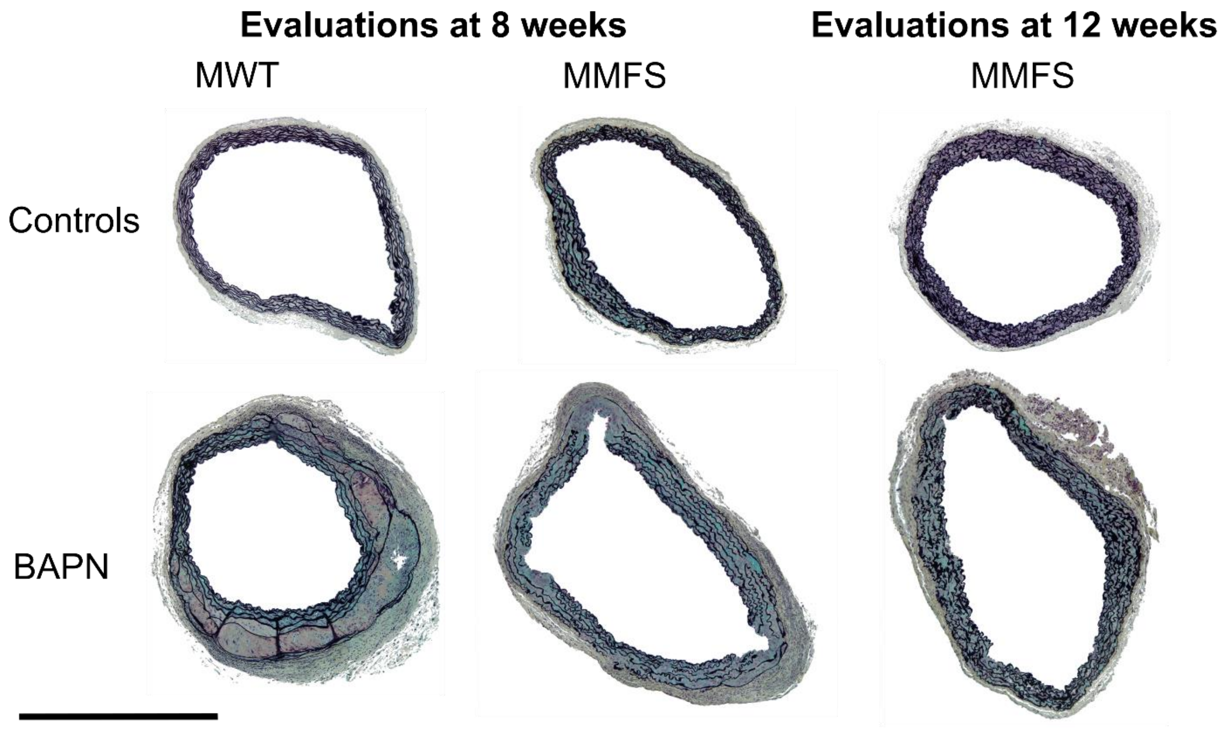


**Figure S5.** Additional Movat-stained histological cross-sections of male (M) wild-type (WT) and *Fbn1*$^{C1041G/+}$ Marfan (MFS) mice with or without 4-week BAPN exposures starting at (left) 4 weeks of age (then evaluated at 8 weeks) or (right) 8 weeks of age (then evaluated at 12 weeks of age), with age- and sex-matched controls not receiving BAPN. These whole sections reveal the spatial heterogeneity and diversity of the phenotype. Note the dissection observed in the MWT aorta following early BAPN exposure (bottom, left) as well as aortic dilatation and adventitial thickening (bottom, middle) in the young MMFS aorta. Scale bar = 1mm.

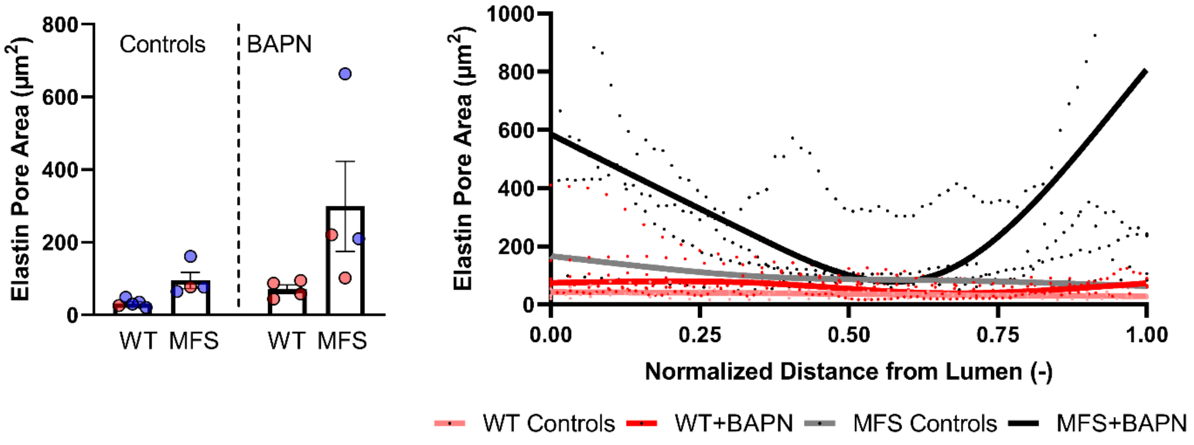


**Figure S6**. Similar to findings for the elastin porosity (Fig 2B-D in the main text), the size of the elastin pores tended to increase in BAPN-exposed groups compared to their matched controls (left panel) and was higher near the endothelial and adventitial layers of the BAPN-exposed MFS group (right panel).

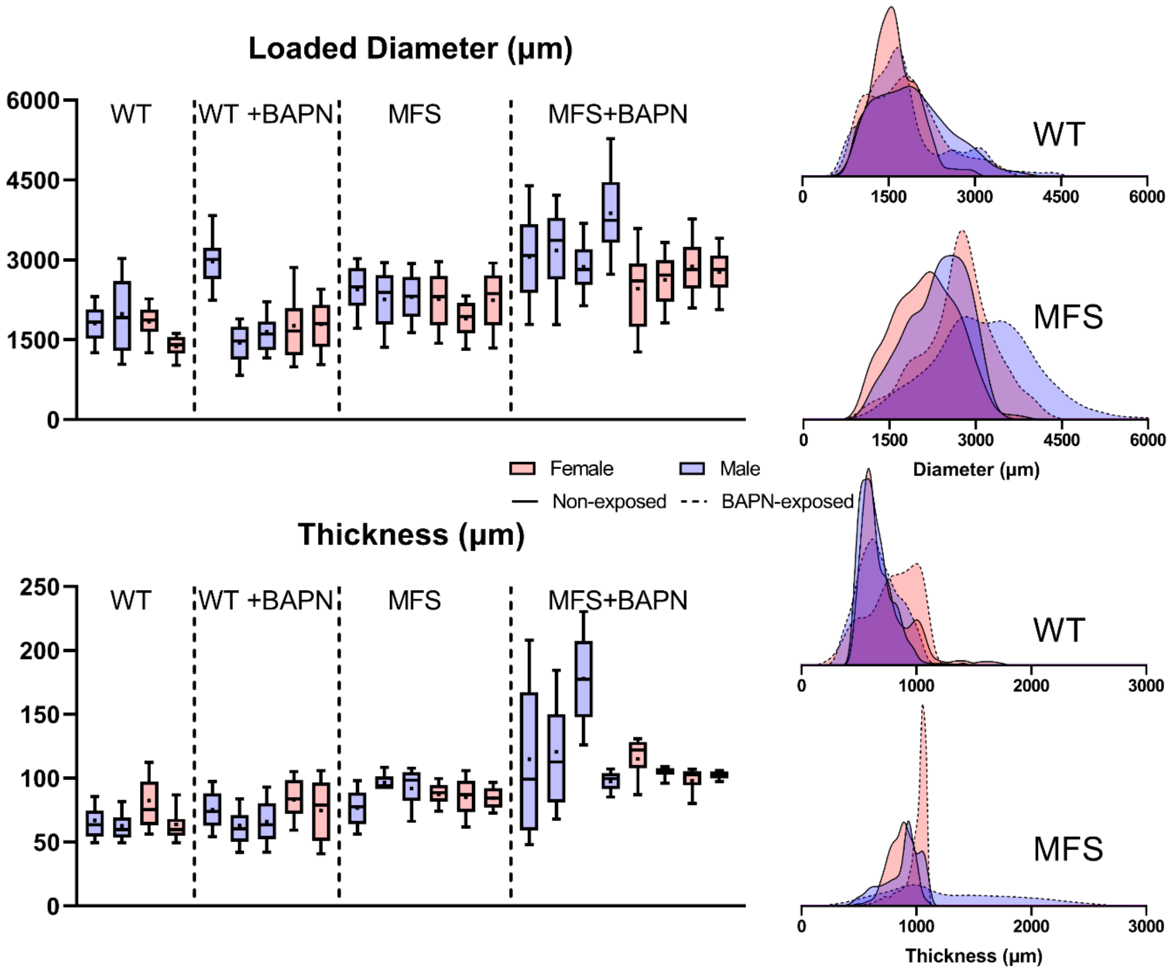


**Figure S7**. Locally identified distributions of outer diameter and thickness for each of the 23 samples tested with the multimodal pDIC+OCT, presented as Box-and-whisker plots (left) and probability density functions (right). The thickness values were measured using OCT and were collected at 100 cross-sections along the length of each vessel and mapped onto the pDIC reconstructed geometry using an automatic co-registration pipeline. Loaded diameter was computed as twice the Euclidean distance from each point on the wall to the nearest point on the curvilinear centerline of the vessel.

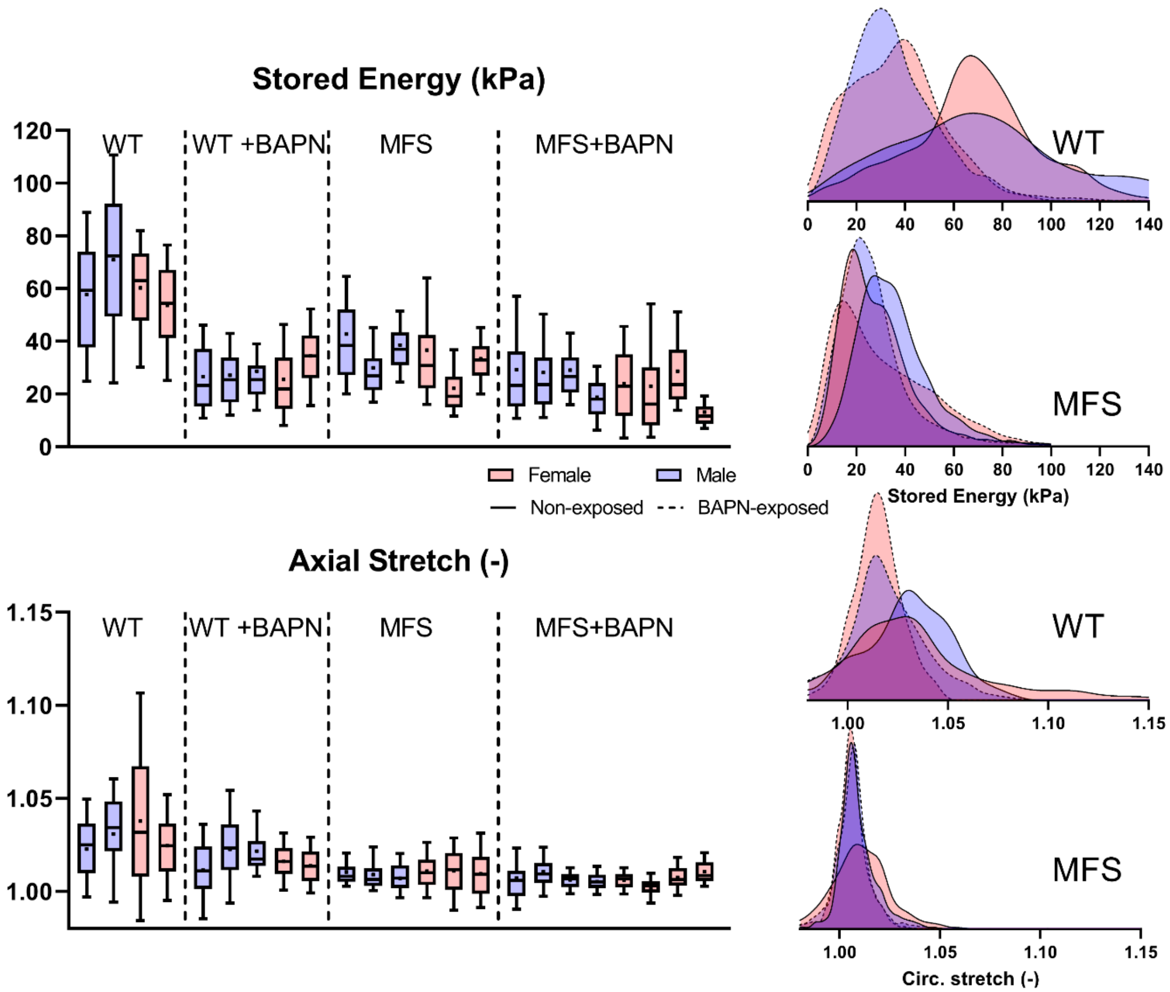


**Figure S8**. Similar to Figure S7, but for elastically stored energy (top row) and (local) axial stretch (bottom row). Values were computed at 100 mmHg relative to the reference configuration (80 mmHg, and *in vivo* axial stretch). MFS, BAPN-exposed MFS, and BAPN-exposed WT mice generally had lower stored energy and axial stretch values compared with WT mice.

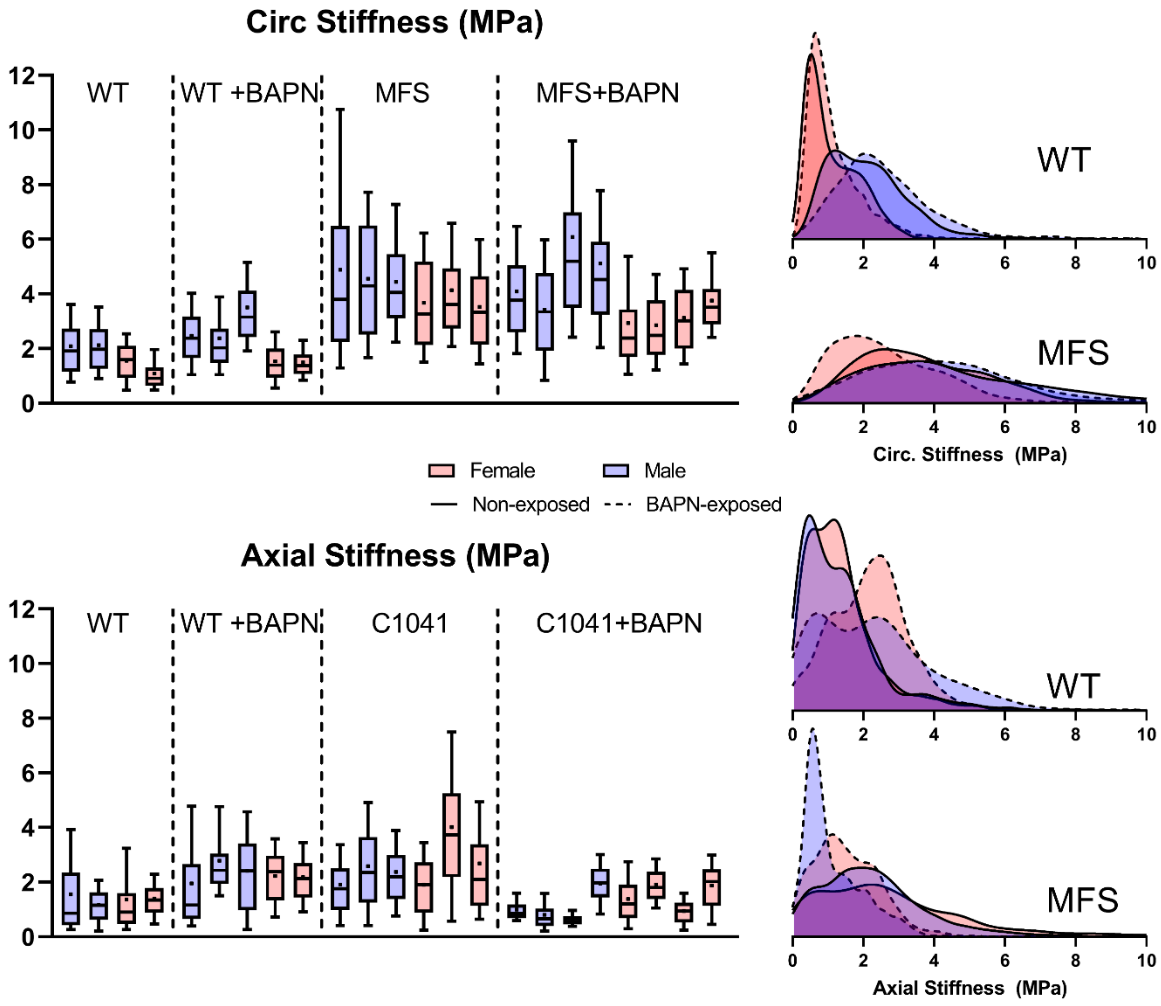


**Figure S9**. Similar to Figs S7-S8, but for specimen-specific circumferential and axial material stiffness. Values were calculated based on local material parameters identified in the inverse characterization but linearized using the theory of small deformations on large (at a common pressure of 100 mmHg and specimen-specific axial stretch). Note the phenotypically increased circumferential stiffness in the Marfan groups that was not affected dramatically by BAPN.

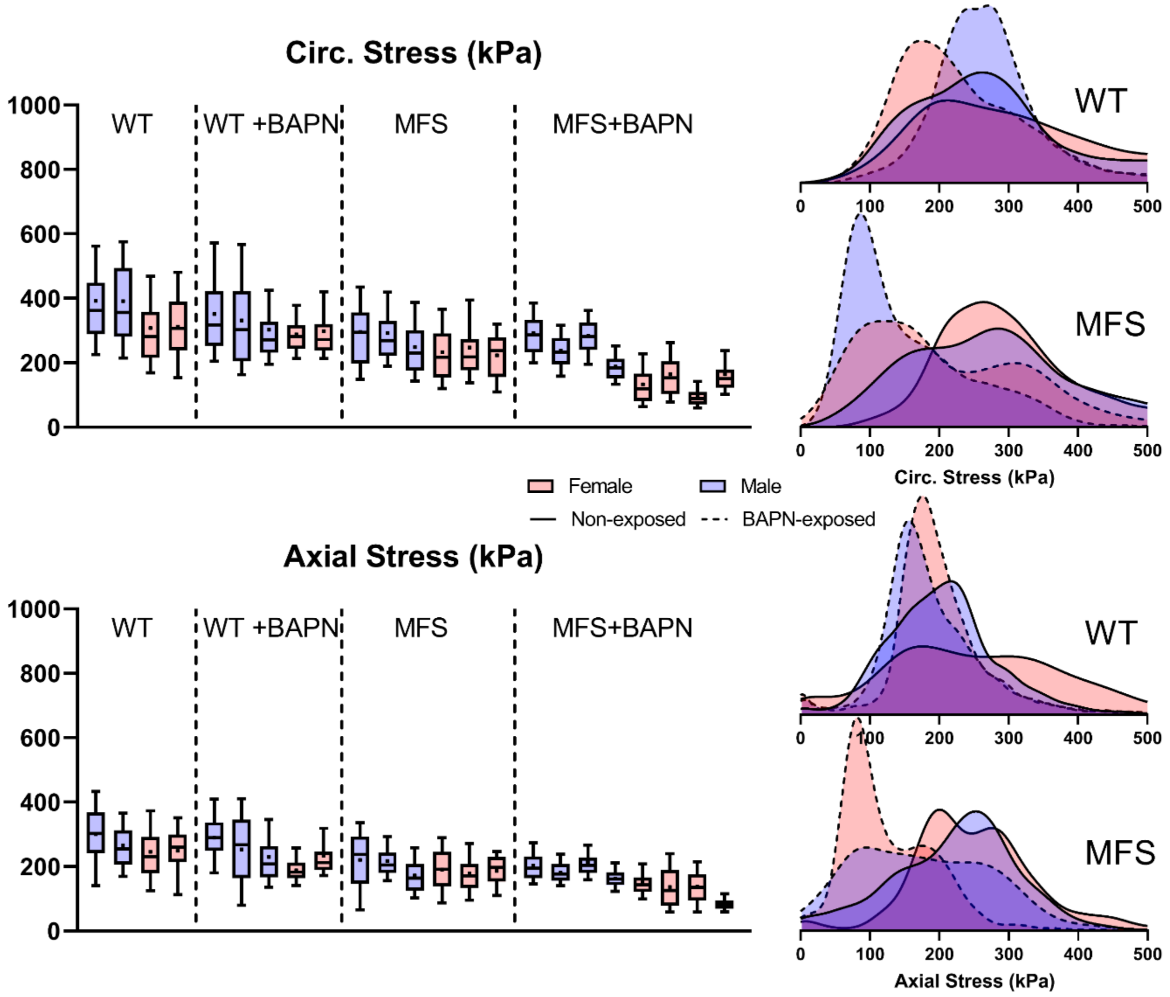


**Figure S10**. Similar to Figs S7-S9, except for circumferential and axial Cauchy stress (kPa). Virtual field-based inverse characterization allowed these values to be computed at any state, shown here for a common pressure of 100 mmHg at the specimen-specific axial stretch. Note the general decrease in both stresses from WT to BAPN-exposed MFS.

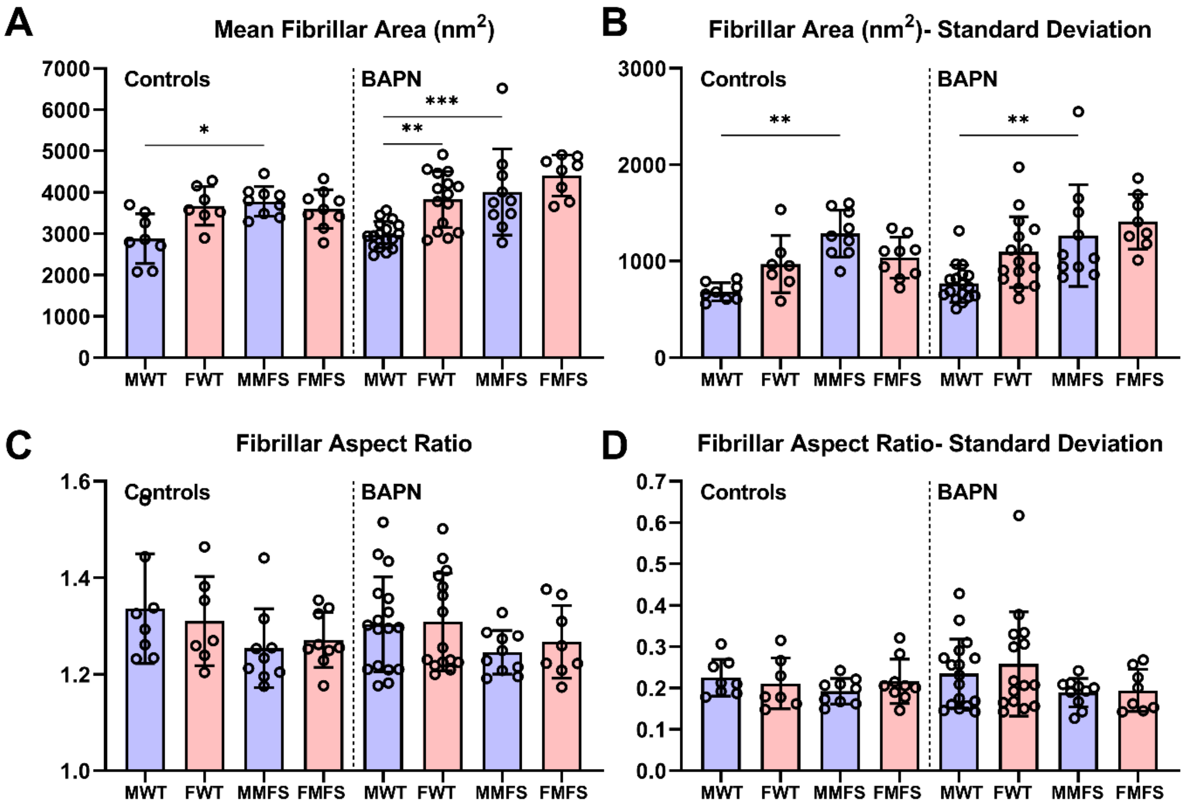


**Figure S11.** In male mice (both without and with 4-week exposures to BAPN), adventitial collagen fibril cross-sectional area was (A) higher and (B) more variable in MFS, on average, compared to age- and sex-matched WT counterparts. Mean fibril area in female WT mice tended to be larger than in male WT mice regardless of BAPN treatment, though this difference was only statistically significant in the BAPN-exposed group due to higher sample sizes. No significant differences were found for fibril aspect ratio (C, D), which was remarkably consistent around 1.3±0.2 across all groups.

**SUPPLEMENTAL FIGURES of METHODS**

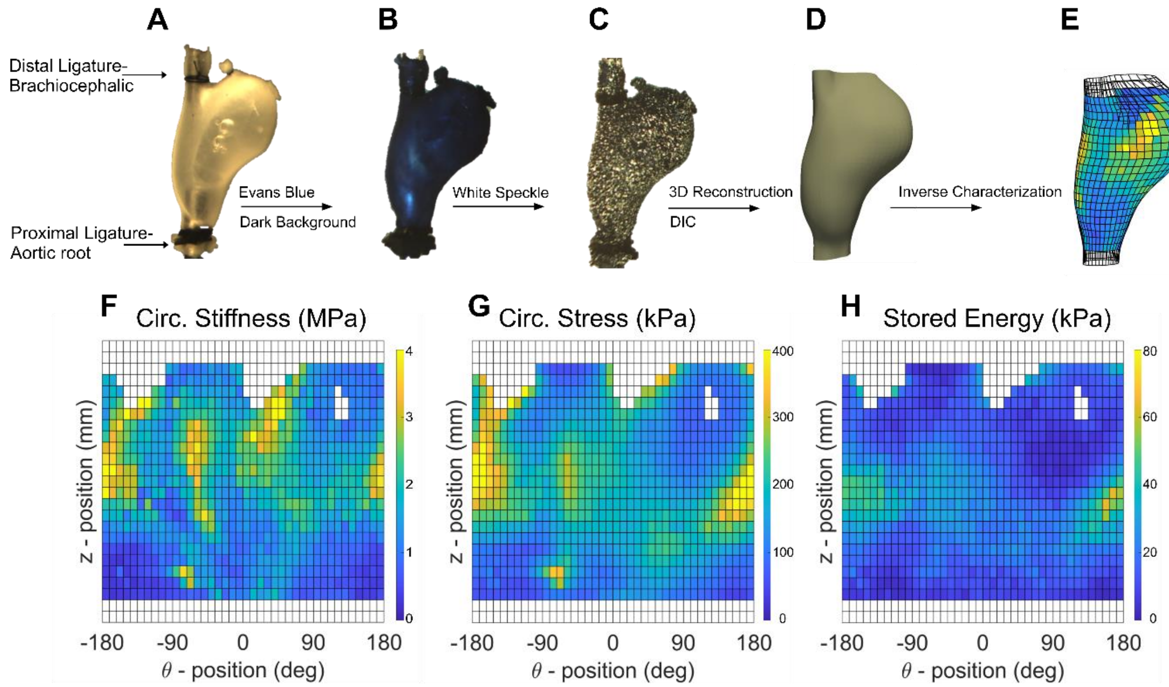


**Figure S12**. Regional material characterization. Following standard biaxial testing a subset of the 8-week old mice, aortas were re-cannulated and secured on a triple blunt-end needle assembly (A), soaked in Evans blue dye to create a dark background (B), and then air-brushed with a white India ink to form a unique speckle pattern on the vessel surface (C). The vessel was then submerged in HBSS-filled 45-degree conical mirror and the reflection of the white pattern was captured by a nearly vertically located digital camera from 8 different rotationally symmetric views at 14 incremental pressures (10-140, 10 mmHg increments) and 3 axial stretches ($\lambda_z^{iv}$ and ±5% of this value) for 336 images per vessel. Digital image correlations over all deformed configurations were then used to reconstruct the 3D surface geometry (D) at each deformed configuration and to compute full-field surface deformations that were used to estimate best-fit parameters of our four-fiber constitutive relation locally at ~1000 elements around the circumference ($\theta$ position) and along the length of each specimen (Z position). The principle of virtual power was enforced at each element to achieve inverse characterization and to compute full-field distributions of the different mechanical metrics, i.e. circumferential stiffness (E, F- unwrapped 2D map), circumferential stress (G), and stored energy (H).

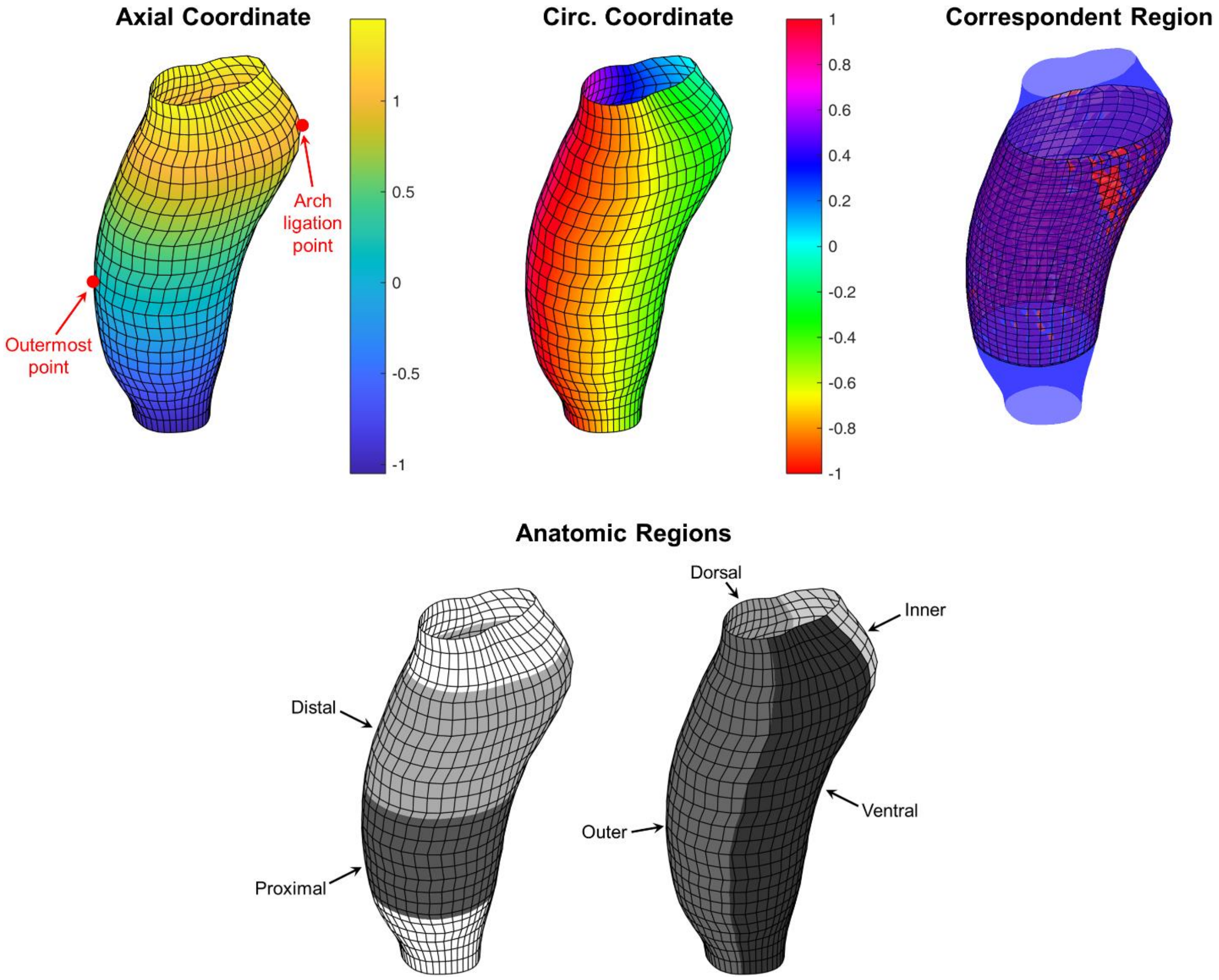


**Figure S13**. Pipeline for parametrization and co-registration of pDIC-based geometric reconstructions, shown here for a representative normal MWT ascending aorta. Along the axial direction, the vessel is parametrized using two anatomic landmarks (red points) such that the axial coordinate is 0 at the outermost point on the outer curvature and 1 at the ligation point on the distal aortic arch (note that parts of the vessel surface lie outside this interval, and thus the axial coordinate values extend beyond the [0, 1] interval). Around the circumference, the vessel is parametrized with periodic boundaries at -1 and 1, located in the middle of the outer curvature. Statistical analyses and modeling were performed only within the "correspondent region" that was spanned by all vessel geometries included in the dataset. For interpretation and presentation of the results, the correspondent region was segmented into two regions along the axial direction (proximal and distal) and four regions circumferentially (inner, dorsal, outer, and ventral).

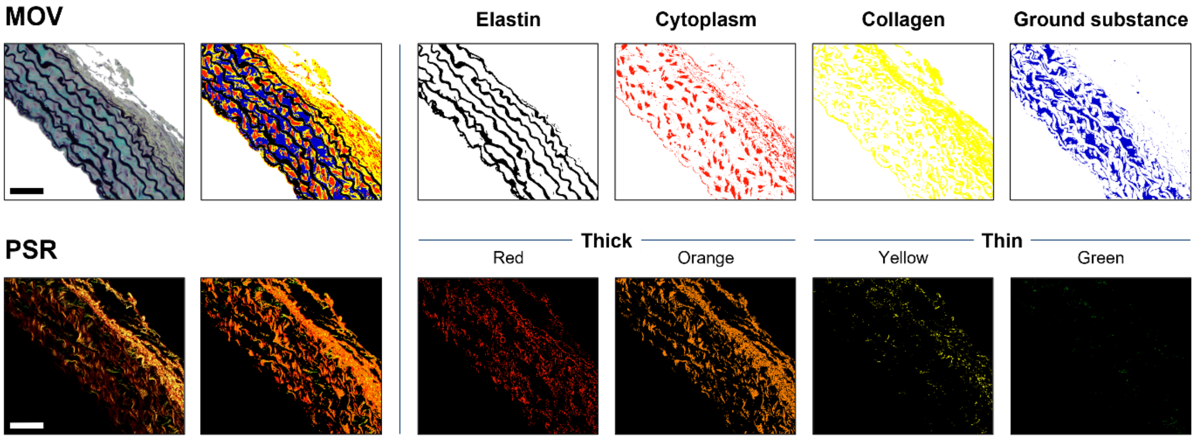


**Figure S14.** Representative histological analysis of (top) Movat (MOV)-stained and (bottom) picro-sirius red (PSR)-stained cross-sections. Color-based analysis routine used to categorize image pixels as elastin (black), cytoplasm (red), fibrillar collagen (yellow), ground substance (blue), or fibrin (now shown) in MOV images, and to detect fibrillar collages in a spectrum from red-to-green in PSR images, digitally separated from the analyzed image for clarity. In PSR images, red/orange and yellow/green pixels were categorized as thick and thin fibers, respectively. MOV image was collected under standard bright-field conditions (180 µs exposure) while PSR image was collected under dark-field conditions using polarized light (32 ms exposure). Scale bar = 100 µm.

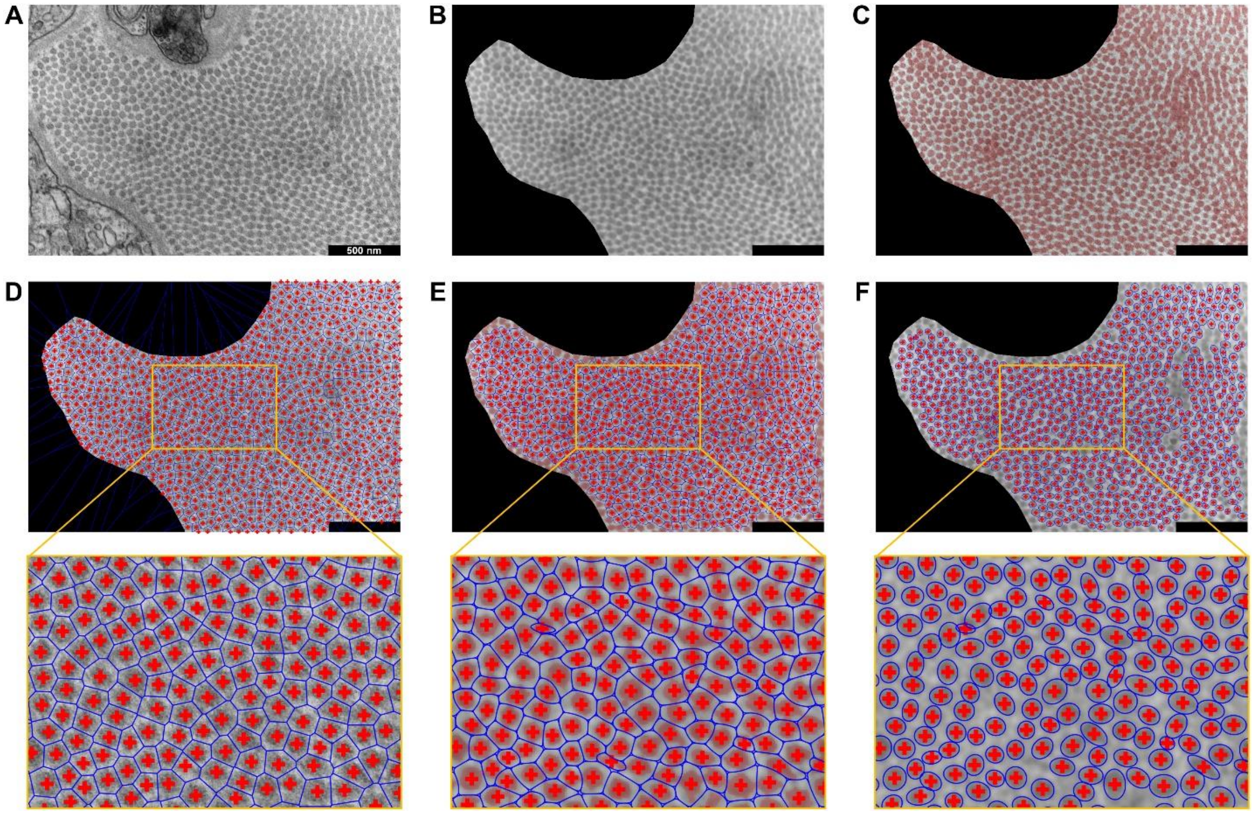


**Figure S15.** Pipeline for semi-automated processing of TEM images to quantify collagen fibril locations, size, and shape. The original acquired image (A) is interactively edited to exclude regions that do not consist of fibrils, smoothed with a Gaussian filter (B), and automatically thresholded as a first attempt to segment fibril and non-fibril pixels (C, with thresholded regions shaded red). Fibril centroid locations (red crosses) are then automatically detected to estimate individual fibril “neighborhoods” as Voronoi cells (D, with neighborhood boundaries shown in blue), which are then interactively corrected by the user if necessary. The user-corrected fibril centroid locations are used to fine-tune the thresholding as a spatially heterogeneous field, and as an initial guess to fit a Gaussian mixture model to the new set of pixels estimated to belong to fibrils (E, where the blue boundary around each fibril neighborhood is the level set at which the membership probability of a thresholded pixel belonging to that fibril equals 50%). Under the approximation that thresholded pixels within a fibril are uniformly distributed within an ellipse, the best-fit ellipse’s size, orientation, and aspect ratio are automatically computed (F).
